# Mu Suppression During Action Observation Only in the Lower, not in the Higher, Frequency Subband

**DOI:** 10.1101/2024.03.28.587150

**Authors:** Ayşe Nur Badakul, Efe Soyman

## Abstract

Mu suppression – desynchronization of neural oscillations in central EEG electrodes during action execution and observation – has been widely accepted as a marker for neural mirroring. It has been conventionally and predominantly quantified in the 8-13 Hz range, corresponding to the alpha frequency band, although few studies reported differences in lower and higher subbands that together constitute the mu frequency band. In the present study, we adopted a data-driven approach to examine the spectral and temporal dynamics of mu suppression when participants watched videos depicting hand and face actions and artificial pattern movements. Our analyses in central EEG electrodes revealed that neural oscillations were significantly suppressed during action observation only in the lower (8-10.5 Hz), not in the higher (10.5-13 Hz), subband. No such subband differentiation was observed for the alpha oscillations in the occipital electrodes. In addition, in the lower subband, significantly stronger suppressions were selective for hand actions in the central EEG electrodes placed over the hand region of the sensorimotor cortices and for facial actions in the frontotemporal electrodes placed over the face region of the sensorimotor cortices. In the higher subband, such stimulus selectivity was only observed for facial actions in the frontotemporal electrodes. Furthermore, the neural oscillations in the lower, but not the higher, subband followed the precise temporal patterning of biological motion in the videos. These results indicate that neural oscillations in the lower subband show the characteristics of neural mirroring processes, whereas those in the higher subband might reflect other mechanisms.

## Introduction

The discovery of mirror neurons in the F5 area of the macaque brain (di Pellegrino et al., 1992; Gallese et al., 1996) has provided a pivotal avenue for unlocking insights into how motor actions are processed in the brain (Rizzolatti & Craighero, 2004). These neurons are active during the observation of others’ actions, in addition to the execution of the same movements, which is widely believed to underlie a mechanism of internal simulation or sensorimotor resonance of observed actions, argued to serve a variety of social cognitive processes (Rizzolatti & Craighero, 2004; Agnew et al., 2007; Cattaneo & Rizzolatti, 2009; Keysers & Gazzola, 2006). Suppression in the power of the mu oscillations recorded from centrally positioned EEG electrodes is similarly seen both during the execution and observation of actions and has thus been considered an EEG marker for this sensorimotor resonance mechanism. First reported as the disappearance of sharp waves, defined as ‘en arceu’, within central electrodes when participants watched actions in a boxing video (Gastaut & Bert, 1954), mu suppression has been conventionally and predominantly quantified in the 8-13 Hz range, corresponding to the alpha frequency band (Muthukumaraswamy & Johnson, 2004; Oberman et al., 2005; Singh et al., 2011; Hobson & Bishop, 2016; Hari et al., 1998; Pineda, 2005; Muthukumaraswamy & Singh, 2008; Aridan et al., 2018), or a slightly narrower or wider frequency band within the 7-16 Hz range (Willemse et al., 2010; Perry & Bentin, 2010; Cheyne et al., 2003; Tamura et al., 2012).

Only a handful of studies examined action observation-induced mu suppression in two discrete subbands – a lower (∼8-10 Hz) and a higher subband (∼10-13 Hz; Cochin et al., 1999; Perry et al., 2010; Frenkel-Toledo et al., 2013; Dumas et al., 2014; Frenkel-Toledo et al., 2014) – suggesting that there might be differences between these subbands regarding functioning of the mirror neuron system (Bazanova & Vernon, 2013; Frenkel-Toledo et al., 2014). Cochin et al. (1999) reported significant mu suppression in the alpha 1 (7.5-10.5 Hz), but not in the alpha 2 (10.5-13 Hz), subband for both observation and execution of finger movements in several electrodes including the central ones. In addition, Perry et al. (2010) showed that intranasal oxytocin significantly enhanced suppression in the low (8-10 Hz), but not in the high (10-12 Hz), subband, during the observation of point-light biological movements in a widely distributed manner. Furthermore, Frenkel-Toledo et al. (2013) also demonstrated significantly stronger action observation-induced mu suppression in the low (8-10 Hz) compared to the high (10-12 Hz) subband. Finally, Dumas et al. (2014) reported significant mu suppression in the lower subband (8-10 Hz) in all central electrodes during action observation in neurotypical participants, whereas, in the upper subband (11-13 Hz), significant suppression was observed in several electrodes across the scalp, but critically not in the central electrodes Cz, C1, C2, C3 and C4, which are most frequently associated with the central mu suppression phenomenon. Despite these differential findings for the two subbands and the more general variability in the selected frequency range for studying mu suppression across studies, a recent meta-analysis did not use a specific frequency range as an inclusion criterion or did not include the frequency range as a moderator in their analyses (Fox et al., 2016). Due to this general neglect of the potential differences between the frequency bands constituting the conventional mu oscillations, several studies still quantify mu suppression within the conventional 8-13 Hz frequency band (e.g., Krol et al., 2020; Lim et al., 2023). We aimed to address these deficits in the literature by adopting a data-driven approach to analyze changes in oscillatory power across a wide range of frequencies during the observation of both biological and non-biological motion.

Mirror neurons were discovered and have been primarily investigated during observation of object-directed hand movements. Since mu suppression has been conceptualized as an indicator of the mirror neuron activity, the same approach was adopted early on, which led to mu suppression being predominantly studied with hand movements encompassing both object-directed hand actions, such as grabbing and reaching (e.g., Muthukumaraswamy et al., 2004; Bernier et al., 2007), as well as non-object-directed hand actions, such as opening and closing the hand (e.g., Oberman et al., 2008; Hobson & Bishop, 2016). Only a limited number of studies examined mu suppression during observation of facial (Karakale et al., 2019), and specifically oral (Muthukumaraswamy et al., 2006; Sakihara & Inagaki, 2015) movements; however, no study to date conducted a systematic comparison of mu suppression responses during observation of different body actions. If mu suppression represents sensorimotor resonance for any internally simulatable biological action, rather than being a mechanism solely responsive to seeing hand movements, mu suppression for different body parts must be seen selectively in the locally specific regions of the sensorimotor cortices where those body parts are represented. Motivated by this rationale, the present study, for the first time in the literature, employed both hand and face movements to investigate mu suppression in different regions of the cortex.

Interestingly, mu suppression has been documented during observation of live actions and dynamic videos showing biological motion (e.g., Wriessnegger et al., 2013; Aridan et al., 2018; Ikeda et al., 2020), as well as static images depicting body parts (e.g., Adolphs et al., 2000; Pineda & Hecht, 2009). Just the simple observation of a static body part, however, cannot solely explain all instances of mu suppression, since further suppression is seen when the observed static body part starts to move (Hobson & Bishop, 2016), suggesting that the observation of the motion itself is a critical factor. To what extent the magnitude of perceived motion is related to the magnitude of mu suppression remains unexamined. To address this gap in our knowledge, the present study assessed, for the first time in the literature, how mu suppression relates to variations in perceived motion, both within the observation of a single action and across observations of different movements.

We addressed these gaps in the literature by examining EEG recordings collected while 30 participants watched videos showing simple hand, face, or artificial pattern movements (hereafter referred to as Hand, Face, and Pattern videos, respectively; Figure 1a: see Supplementary Table 1 for the full list of hand and face movements). In parallel with Hobson and Bishop (2016), each video started with a 4-second static period, followed by a 2-second dynamic period depicting the movement, and finally a 2-second static post-stimulus period. Participants had to rate the perceived motion in a random quarter of all trials, which was implemented to motivate sustained attention to the videos. We used several different movements with varying magnitudes of motion within each stimulus category to assess the relationships between perceived motion and neural responses (Figure 1b). In addition, the same movement was repeated twice in each video to investigate repetition-dependent temporal dynamics in neural responses. Suppression of neural activity was quantified by taking the last 2 seconds of the initial 4-second static period as the baseline and the 2-second dynamic period as the active period, in line with Hobson and Bishop (2016). We primarily focused on the average of C3 and C4, T7 and T8, and O1 and O2 electrodes (hereafter referred to as Central, Temporal, and Occipital sites, respectively), since C3 and C4 electrodes overlay the hand area and T7 and T8 electrodes overlay the face area of the sensorimotor cortices. O1 and O2 electrodes were analyzed to control for alpha activity originating in the visual cortex (Hobson & Bishop, 2016; Hobson & Bishop, 2017). After the EEG recordings, participants completed the Interpersonal Reactivity Index (Davis, 1983) to assess whether inter-individual differences in mu suppression are related to empathic skills, for which the findings are conflicting in the literature (Cheng et al., 2008; Yang et al., 2009; Woodruff et al., 2011; DiGirolamo et al., 2019).

**Figure 1.**
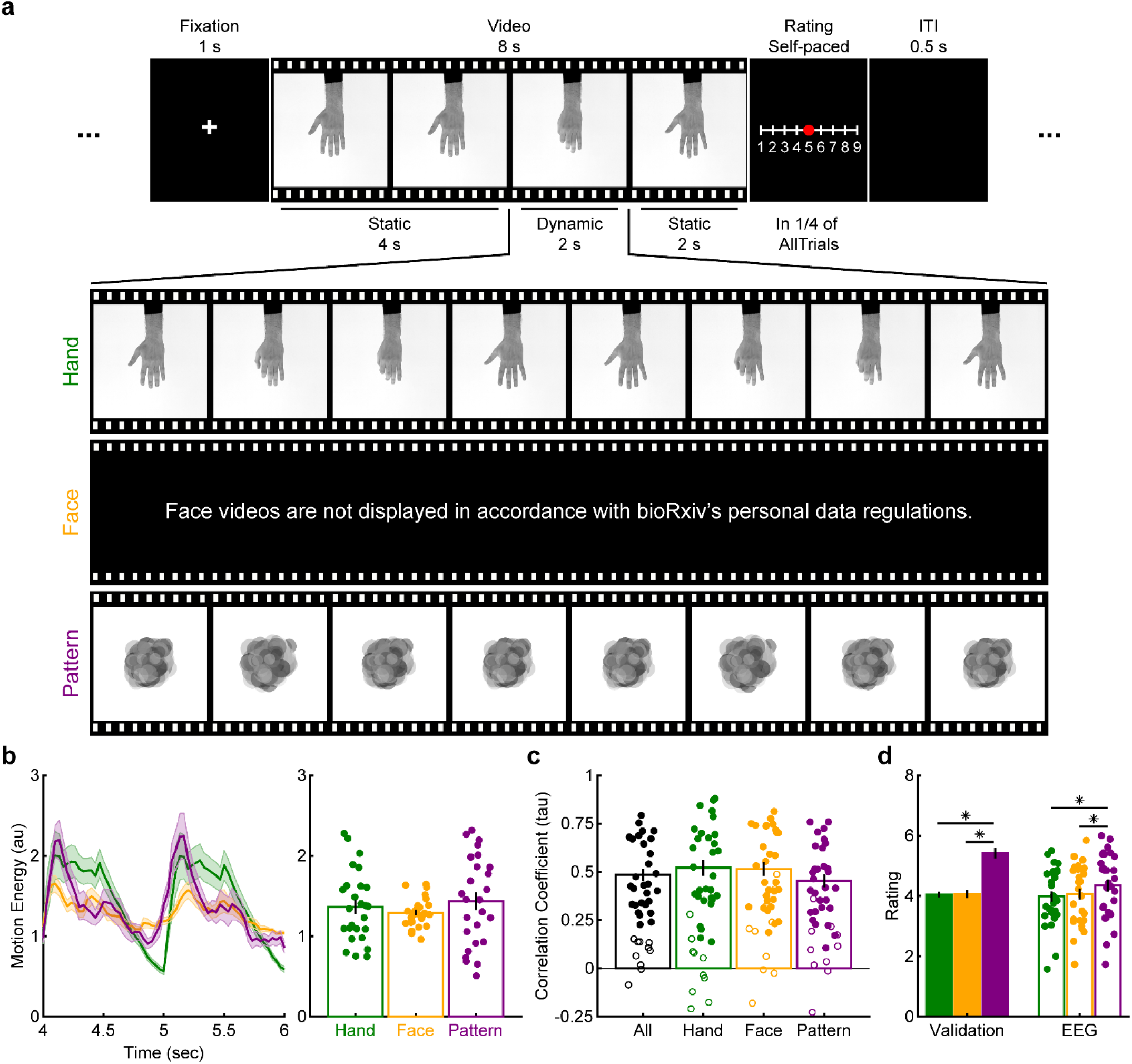
Stimuli, experimental design, and motion ratings. **(a)** Each trial started with a 1-second fixation point, followed by an 8-second video, the first 4 seconds of which displayed a static hand or face in its neutral state or an artificially created pattern. The static image turned into a dynamic video displaying different magnitudes of motion in the next 2 seconds, which was followed by another 2-second static post-motion period. In a pseudo-randomly determined quarter of the trials, participants were asked to rate the perceived motion level in the video. The dynamic portion of the videos displayed simple hand or face actions (see Supplementary Table 1 for the full list of movements), or random movements of an artificial pattern. For all videos, the same movement was repeated twice within 2 seconds. **(b)** Mean time-resolved (left) and time-averaged (right) motion energy values in Hand, Face, and Pattern videos. **(c)** Mean correlation coefficients between average motion ratings in a separate validation study and the motion ratings of the participants in the present study. Filled circles show the correlation coefficients of 30 participants included in further statistical analyses, whereas open circles represent the correlation coefficients of 10 excluded participants. **(d)** Mean motion ratings for Hand, Face, and Pattern videos in the validation and the present study. For all analyses, shadings and error bars depict SEMs. For analyses in (d), asterisks indicate significant differences between pairs of stimulus categories, and the significance level was Bonferroni-corrected at 0.05/3=0.017.

## Results

### Perceived Motion Ratings During EEG Recordings are in Line with Normative Ratings

To assess whether the participants attended the video stimuli and engaged in the motion rating task during EEG recordings, we first analyzed participants’ ratings in relation to those of a separate sample of 41 participants, who rated each video in an earlier validation study. The correlation coefficients between the participants’ own motion ratings and the average validation ratings across all stimuli were significantly positive for 30 participants, who were thus determined to have sufficiently attended the videos during EEG recordings (Figure 1c). The remaining 10 participants who failed to show significant correlations were excluded from all further analyses. The correlation coefficients between participant ratings and the validation ratings did not significantly differ across Hand, Face, and Pattern videos (*F_2,58_*=3.401, *p*=0.040, *BF_inc_*=1.345; planned pairwise comparisons did not reach statistical significance at the Bonferroni-corrected alpha value of 0.05/3=0.017; Figure 1c). Average motion ratings, on the other hand, were significantly greater for Pattern videos (*F_2,58_*=8.303, *p*=7×10^-4^, *BF_inc_*=42.381) compared to Hand (*t_29_*=4.156, *p*=3×10^-4^, *BF_10_*=109.212) and Face videos (*t_29_*=2.607, *p*=0.014, *BF_10_*=3.334), with no difference between Hand and Face videos (*t_29_*=0.988, *p*=0.331, *BF_10_*=0.304; Figure 1d). The same pattern of results was found in the validation sample (*F_1.75,69.86_*=59.759, *p*=9×10^-15^, *BF_inc_*=9×10^13^; Pattern vs Hand: *t_40_*=9.372, *p*=10^-11^, *BF_10_*=8×10^8^; Pattern vs Face: *t_40_*=8.218, *p*=4×10^-10^, *BF_10_*=3×10^7^; Hand vs Face: *t_40_*=0.124, *p*=0.902, *BF_10_*=0.170; Figure 1d). These findings suggest that the 30 participants included in the statistical analyses of EEG recordings attended the video stimuli and rated perceived motion similarly to an independent validation sample.

### Low and High Subbands Differ in Degrees of Suppression, Stimulus-Selectivity, and Temporal Dynamics

We analyzed the EEG recordings by utilizing a cluster-correction algorithm to examine time-frequency points that showed significant modulation with motion perception. Figure 2a depicts clusters with significant enhancement or suppression of neural activity from the 2-second static baseline period of videos to the 2-second active period for Hand, Face, and Pattern videos in Central, Temporal, and Occipital sites (see Supplementary Figures 1-3 for all frequencies within the 0-30 Hz range for all recording sites). Whereas Occipital sites revealed a strong and homogenous suppression in neural activity starting with the onset of motion for all stimulus categories in frequencies extending the conventional alpha band of 8-13 Hz, Central sites indicated very different patterns for frequencies below and above approximately the middle of this frequency band. That is, there was a significant suppression in neural responses below ∼10.5 Hz for all stimulus categories (Hand: 10.75 Hz; Face: 10 Hz; Pattern: 10.25 Hz); while above ∼10.5 Hz, there was a significant increase in neural activity for Pattern videos and no significant change for Hand and Face videos (Figure 2a). Temporal sites revealed an in-between pattern, indicating significant suppression of neural activity homogeneously in the 8-13 Hz frequency band for Face videos, while the suppression was limited to ∼≤11 Hz for Hand and Pattern videos (Figure 2a).

**Figure 2.**
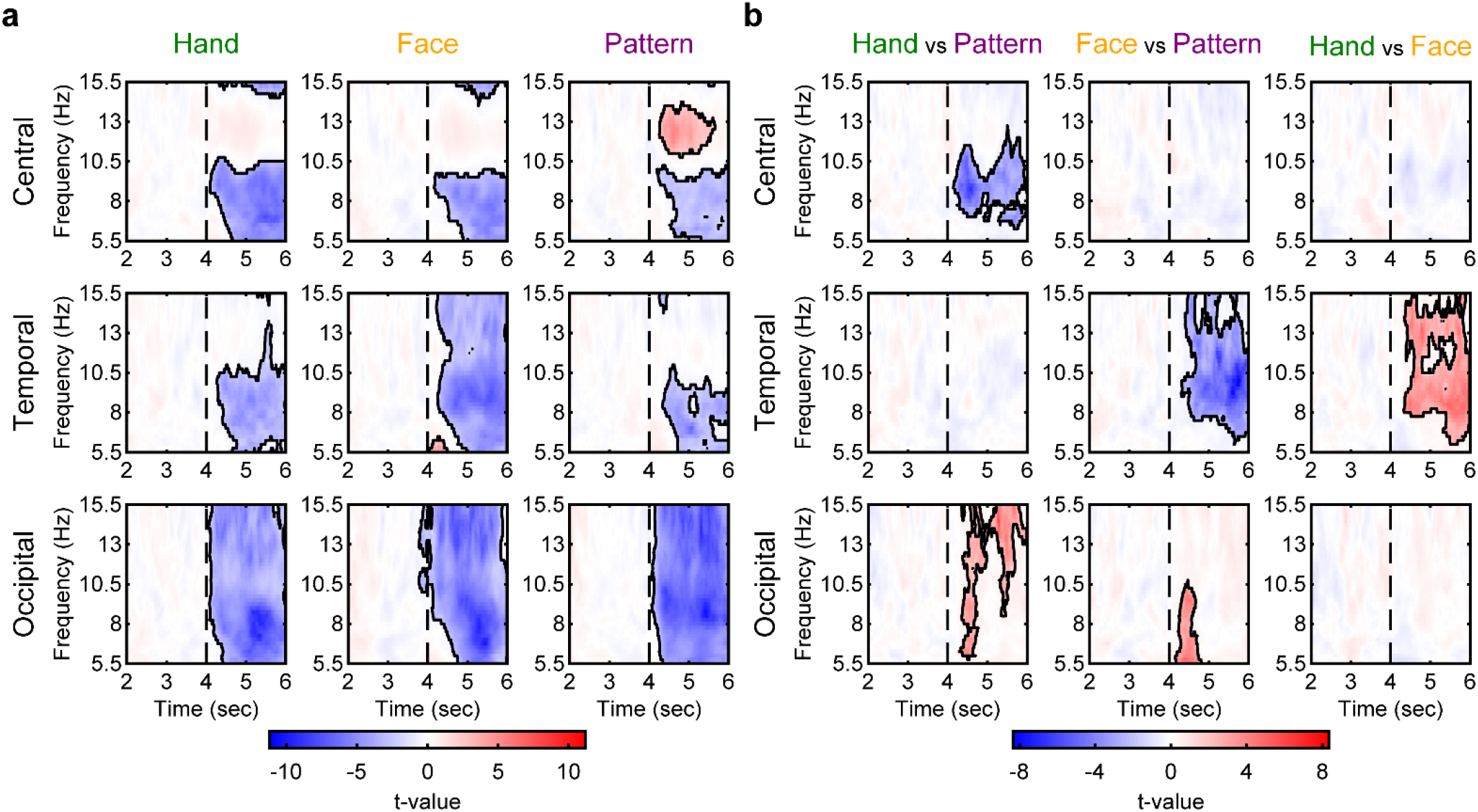
Time-frequency representations of neural responses. **(a)** T-values of one-sample t-tests comparing log power change values to zero across frequencies and time points for Hand, Face, and Pattern videos in Central, Temporal, and Occipital sites. **(b)** T-values of paired-sample t-tests comparing log power change values between pairs of stimulus categories across frequencies and time points for Hand, Face, and Pattern videos in Central, Temporal, and Occipital sites. For all analyses, the sample size was 30, and statistical significance was determined via cluster correction. Statistically significant clusters are shown with black contours and non-significant t-values are scaled by a factor of 0.25 for better visualization of the significant clusters. For analyses in (b), colder colors indicate smaller log power change values for the first stimulus category in the title of the plot, whereas warmer colors indicate smaller values for the second stimulus category. Note that, during the initial 2 seconds, the video was static, and during the ladder 2 seconds, the video displayed motion.

We next employed a similar cluster-correction method to examine the time-frequency points that showed significant differences between pairs of stimulus categories. In Central sites, there was significantly more suppression only for Hand compared to Pattern videos, concentrated in frequencies from ∼7 to ∼11 Hz (Figure 2b). Temporal sites revealed significantly stronger suppression for Face videos compared to both Hand and Pattern videos within the 8-13 Hz frequency band relatively homogenously (Figure 2b). Finally, Occipital sites demonstrated significantly more suppression of neural activity for Pattern videos compared to Hand and Face videos within the alpha frequency band, although these significant clusters were smaller and more variable in frequency and time than the significant clusters for Hand and Face videos (Figure 2b).

Taken together, these results suggest that the suppression of mu oscillations in Central sites during observation of biological motion, which is typically presumed to be a homogenous effect across the 8-13 frequency band, does, in fact, consist of two frequency subbands that show differential patterns: a lower frequency band (8-10.5 Hz; hereafter referred to as Low subband) that indeed shows motion perception-dependent suppression more strongly for hand movements compared to non-biological motion, and a higher frequency band (10.5-13 Hz; hereafter referred to as High subband) that does not show suppression and does not discriminate between biological and non-biological motion. Subsequent analyses were performed by considering the neural responses in these two subbands separately to further investigate their temporal and stimulus-selectivity characteristics.

Figure 3 depicts the temporal profiles of neural responses in the Low (8-10.5 Hz) and High (10.5-13 Hz) subbands separately. As seen before, in Central sites, neural responses showed opposing trends of suppression and enhancement in Low and High subbands, respectively. Only Low subband responses showed significantly more suppression for Hand than Pattern videos in an early time window between 4.27-4.81 sec and in a shorter later time window between 5.38-5.52 sec (Figure 3a). In Temporal sites, Face videos elicited stronger suppression compared to Pattern videos in both Low and High subbands starting from 4.57-5.99 sec onwards (Figure 3a,b). Finally, Occipital sites revealed similarly suppressed responses for all stimulus categories in both Low and High subbands (Figure 3a,b). Overall, these findings corroborate earlier analyses by demonstrating different patterns for Low and High subbands in Central sites and add to them by revealing temporal differences between Hand responses in Central sites and Face responses in Temporal sites. Subsequent analyses were performed by analyzing neural responses in the earlier (4-5 seconds; hereafter referred to as the Early period) and the ladder (5-6 seconds; hereafter referred to as the Late period) periods separately to further probe these temporal differences in detail.

**Figure 3.**
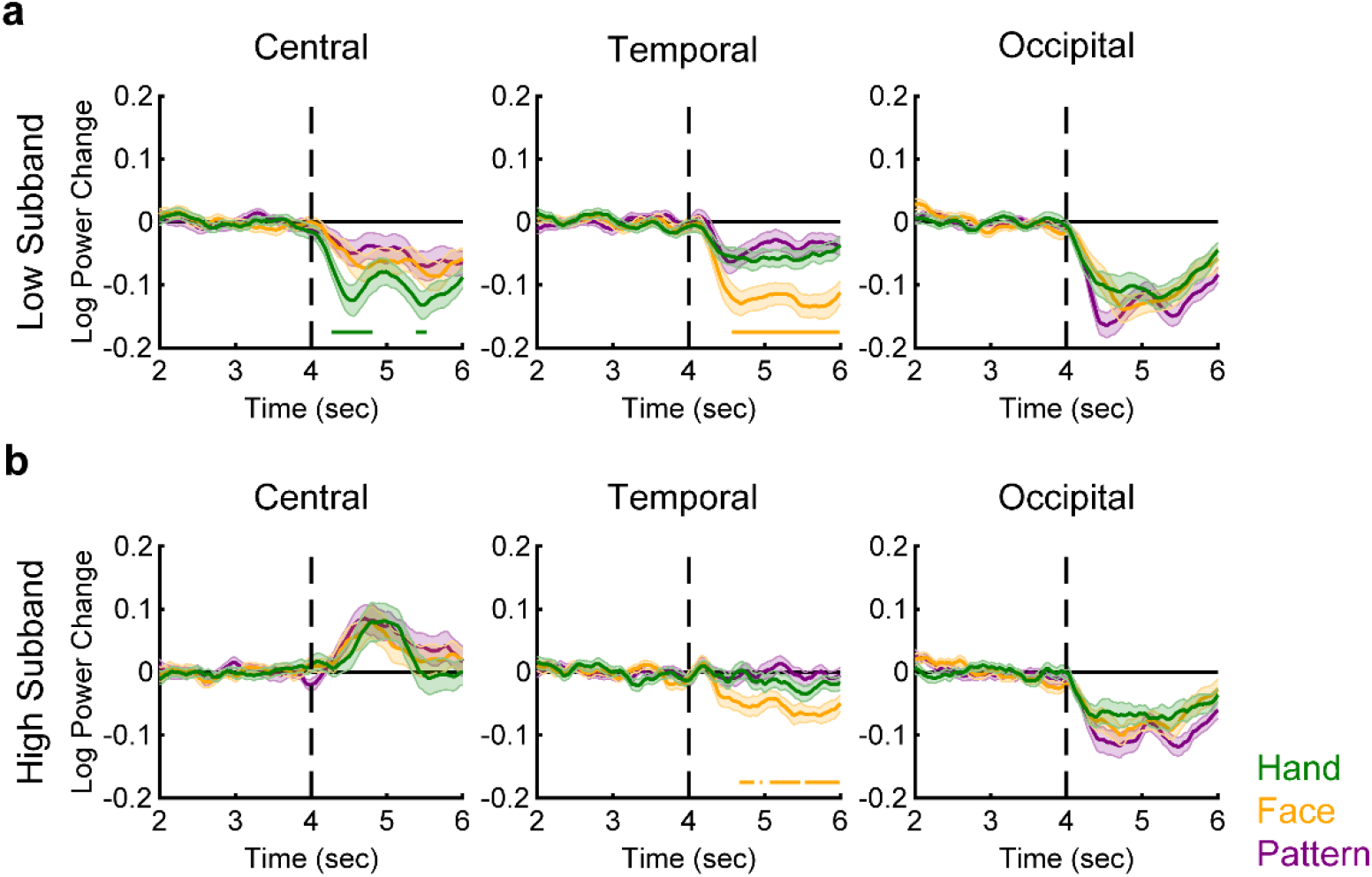
Temporal dynamics of Low and High subband responses. Mean log power change values in **(a)** Low and **(b)** High subbands as a function of time for Hand, Face, and Pattern videos in Central, Temporal, and Occipital sites. Shadings depict SEMs. Horizontal bars within individual plots indicate time windows during which Hand (green) or Face (orange) responses were significantly lower than Pattern responses. For all analyses, the sample size was 30 and the significance level was corrected for false discovery rate at a *q* level of 0.05. Note that, during the initial 2 seconds, the video was static, and during the ladder 2 seconds, the video displayed motion.

A 2 subband (Low, High) x 2 period (Early, Late) x 3 site (Central, Temporal, Occipital) x 3 stimulus (Hand, Face, Pattern) repeated-measures ANOVA showed that both subband and period factors significantly interacted with other factors (subband x site: *F_1.44,41.61_*=13.463, *p*=2×10^-4^, *BF_inc_*=3×10^4^; subband x site x stimulus: *F_3.02,87.47_*=5.094, *p*=0.003, *BF_inc_*=143.562; period x site x stimulus: *F_4,116_*=2.894, *p*=0.025, *BF_inc_*=1.010). Thus, the two subbands and the two periods were further investigated via four separate 3 site (Central, Temporal, Occipital) x 3 stimulus (Hand, Face, Pattern) repeated-measures ANOVAs. In all four of these ANOVAs, there were significant interactions between site and stimulus (*F_4,116_*≥4.018, *p*≤0.004, *BF_inc_*≥4.860, Figure 4). Significant suppression was observed for all stimuli in all recording sites (*t_29_*≥3.092, *p*≤0.004, *BF_10_*≥9.151; see Supplementary Table 2 for all statistics), except for Pattern videos in Central and Temporal sites. Low subband responses in the Early period revealed significantly more suppression for Hand compared to Face and Pattern videos in Central sites and significantly more suppression for Face compared to Hand and Pattern videos in Temporal sites (*t_29_*≥3.212, *p*≤0.003, *BF_10_*≥11.899; Figure 4a; see Supplementary Table 3 for all statistics). Low subband responses in the Late period showed exactly the same pattern of results in terms of both significant differences between the stimulus categories (*t_29_*≥3.149, *p*≤0.004, *BF_10_*≥10.360; Figure 4a) and significant suppressions (*t_29_*≥3.608, *p*≤0.001, *BF_10_*≥29.356). Despite the significant interaction between site and stimulus, High subband responses in the Early period did not reveal any significant stimulus difference in any of the recording sites (*t_29_*≤2.765, *p*≥0.010, *BF_10_*≤4.583, not statistically significant at the Bonferroni-corrected alpha level of 0.05/9=0.0056, Figure 4b). There was a significant enhancement of High subband responses in this Early period for Pattern videos in Central sites (*t_29_*=3.235, *p*=0.003, *BF_10_*=12.539), while in Occipital sites, there was a significant suppression for all stimulus categories (*t_29_*≥4.338, *p*≤2×10^-4^, *BF_10_*≥171.274, Figure 4b). Finally, High subband responses in Temporal sites during the Late period were significantly more suppressed for Face than for Hand and Pattern videos (*t_29_*≥3.397, *p*≤0.002, *BF_10_*≥18.042, Figure 4b). There were significant suppressions for Face videos in Temporal sites and for all stimulus categories in Occipital sites (*t_29_*≥4.443, *p*≤10^-4^, *BF_10_*≥222.126, Figure 4b). To sum up, stronger mu suppression during the observation of hand actions compared to non-biological motion was only seen in the lower frequency subband and relatively soon after the initiation of the movement, whereas stronger mu suppression during the observation of facial actions compared to non-biological motion was seen in both the Low and the High subbands in a relatively later time window.

**Figure 4.**
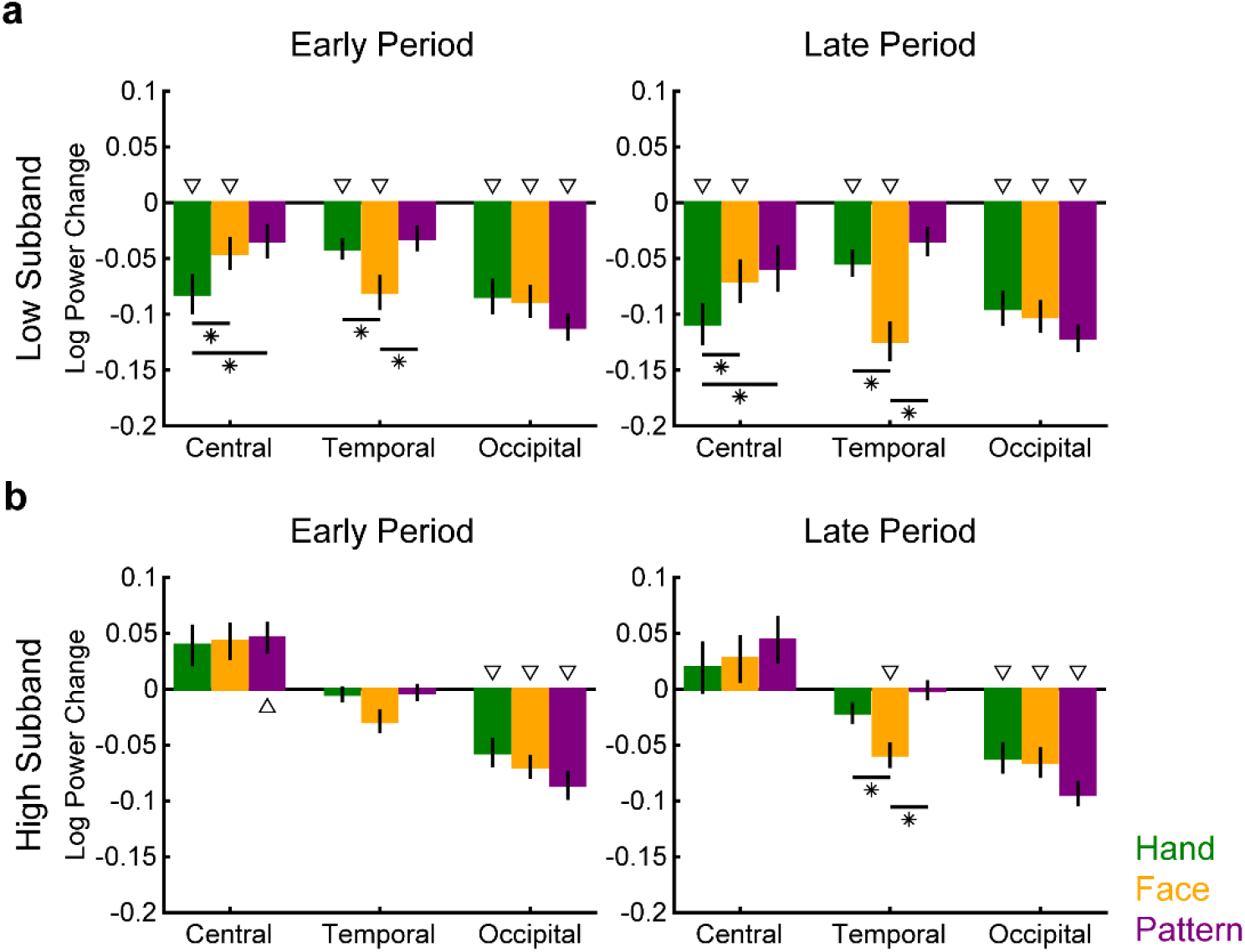
Low and High subband responses in Early and Late periods. Mean log power change values in **(a)** Low and **(b)** High subbands in Early (left) and Late (right) periods for Hand, Face, and Pattern videos in Central, Temporal, and Occipital sites. Error bars depict SEMs. Asterisks indicate significant differences between pairs of stimulus categories. Triangles indicate significant differences from zero. For all analyses, the sample size was 30 and the significance level was Bonferroni-corrected at 0.05/9=0.006 (see Supplementary Tables 2 and 3 for all statistics).

We also conducted a more comprehensive analysis of the Low and High subband responses in Early and Late periods in all 32 recording sites. Low subband responses indicated significant suppressions during both Early and Late periods in the great majority of the electrodes for Hand and Face videos, whereas significant suppressions were limited largely to the occipital and parietal sites for Pattern videos (Figure 5a-c; see Supplementary Table 4 for the full list of significant recording sites). On the other hand, significant suppressions in High subband responses during both Early and Late periods were mainly limited to the occipital and parietal recording sites for all stimulus categories, except some lateral temporal, frontal, and frontocentral sites for Face videos, which also revealed significant suppressions with motion perception (Figure 5a-c). Pairwise comparisons of stimulus categories revealed very few sites with significant differences. Significantly more suppression for Hand compared to Pattern videos was only observed in Low subband responses during the Early period in C3, Cz, C4, and Fp1 electrodes (Figure 5d). Both Low and High subband responses were significantly more suppressed for Face compared to Pattern videos only during the Late period in lateral temporal, frontal, and frontocentral sites (Figure 5e, see Supplementary Table 5 for the full list of significant recording sites). These results support the above findings in a spatially more detailed topography.

**Figure 5.**
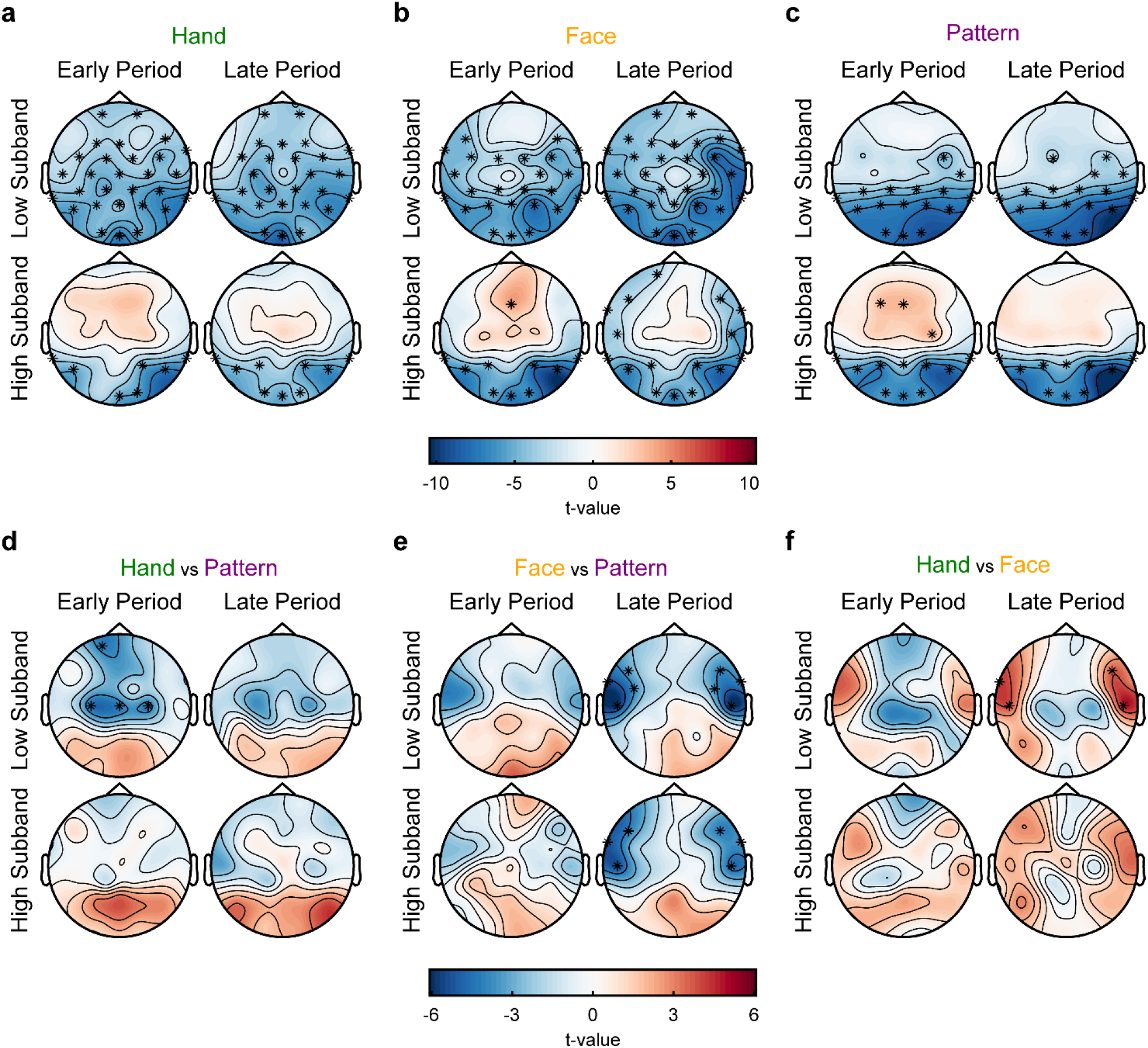
Topographical representations of Low and High subband responses in Early and Late periods. T-values of one-sample t-tests comparing log power change values to zero in Low and High subbands in Early and Late periods for **(a)** Hand, **(b)** Face, and **(c)** Pattern videos. T-values of paired-samples t-tests comparing **(d)** Hand and Pattern, **(e)** Face and Pattern, and **(f)** Hand and Face responses in Low and High subbands in Early and Late periods. For all analyses, the sample size was 30 and the significance level was corrected for false discovery rate at a *q* level of 0.05. For analyses in (a-c), asterisks indicate significant differences from zero. For analyses in (d-f), asterisks indicate significant differences between pairs of stimulus categories (see Supplementary Tables 4 and 5 for the full list of significant recording sites). For analyses in (d-f), colder colors indicate smaller log power change values for the first stimulus category in the title of the plot, whereas warmer colors indicate smaller values for the second stimulus category.

### Re-Analysis of a Pre-Registered Report Supports Our Main Findings Regarding Low and High Subband Differentiation

The video stimuli we used in the present study were largely motivated by, thus in many respects similar to, the videos used in the pre-registered report by Hobson and Bishop (2016). Videos in this study depicted hand movements without any objects (HNO), the same movements with an object in hand (HO), or kaleidoscopic motion as control stimuli. Central mu responses in Hobson and Bishop (2016) were analyzed via directly averaging across the 8-13 Hz frequency band; thus, whether there were differences in Low and High subband responses similar to what we observed in our analyses were unexplored. We took advantage of the fact that the raw data files and the analysis scripts were shared by the authors (Hobson, 2015; https://osf.io/yajkz/) and re-analyzed the dataset using our analysis pipelines. Since there were no Face videos in the Hobson and Bishop (2016) study, Temporal sites were not included in these analyses.

A 2 subband (Low, High) x 2 period (Early, Late) x 2 site (Central, Occipital) x 3 stimulus (HNO, HO, Kaleidoscope) repeated-measures ANOVA showed that subband and period significantly interacted with other factors (subband x site: *F_1,60_*=45.876, *p*=6×10^-9^, *BF_inc_*=4×10^9^; subband x period: *F_1,60_*=12.073, *p*=10^-5^, *BF_inc_*=5×10^4^; period x site: *F_1,60_*=4.355, *p*=0.041, *BF_inc_*=6703.152; period x stimulus: *F_2,120_*=5.955, *p*=0.003, *BF_inc_*=63.192; subband x period x site: *F_1,60_*=20.828, *p*=3×10^-5^, *BF_inc_*=2×10^4^). Thus, the two subbands and the two periods were further analyzed via four separate 2 site (Central, Occipital) x 3 stimulus (HNO, HO, Kaleidoscope) repeated-measures ANOVAs. These four ANOVAs revealed significant site x stimulus interactions (*F_2,120_*≥6.170, *p*≤0.003, *BF_inc_*≥15.708). Low subband responses during the Early period were significantly suppressed with motion perception for only HNO and HO videos in Central sites and for all stimulus categories in Occipital sites (*t_60_*≥2.943, *p*≤0.005, *BF_10_*≥6.829, Figure 6a; see Supplementary Table 6 for all statistics). In Occipital sites, suppression was stronger for Kaleidoscope videos compared to HNO and HO videos (*t_60_*≥3.495, *p*≤9×10^-4^, *BF_10_*≥29.362, Figure 6a; see Supplementary Table 7 for all statistics). Notably, there was a strong, but non-significant trend for more suppression for HNO than Kaleidoscope videos in Central sites (*t_60_*=2.072, *p*=0.043, *BF_10_*=1.026, not significant at the Bonferroni-corrected alpha level of 0.05/6=0.0083, Figure 6a). Low subband responses during the Late period were significantly suppressed for all stimulus categories in both Central and Occipital sites (*t_60_*≥2.854, *p*≤0.006, *BF_10_*≥5.499) and were significantly more suppressed for Kaleidoscope videos compared to HNO and HO videos in Occipital sites (*t_60_*≥4.774, *p*≤10^-5^, *BF_10_*≥1535.484, Figure 6a). High subband responses in Occipital sites were, in both Early and Late periods, significantly lower for Kaleidoscope compared to HNO and HO videos (*t_60_*≥3.223, *p*≤0.002, *BF_10_*≥14.000, Figure 6b). Whereas High subband responses were significantly suppressed with motion perception in Occipital sites for all stimulus categories during both Early and Late periods (*t_60_*≥4.781, *p*≤10^-5^, *BF_10_*≥1574.756), there was a significant enhancement of High subband responses in Central sites only for Kaleidoscope videos during the Early period (*t_60_*=3.025, *p*=0.004, *BF_10_*=8.393, Figure 6b). These results are largely in line with our findings in Central electrodes indicating significant mu suppression only in the lower frequency subband, while significant enhancement was seen for non-biological motion in the higher frequency subband.

**Figure 6.**
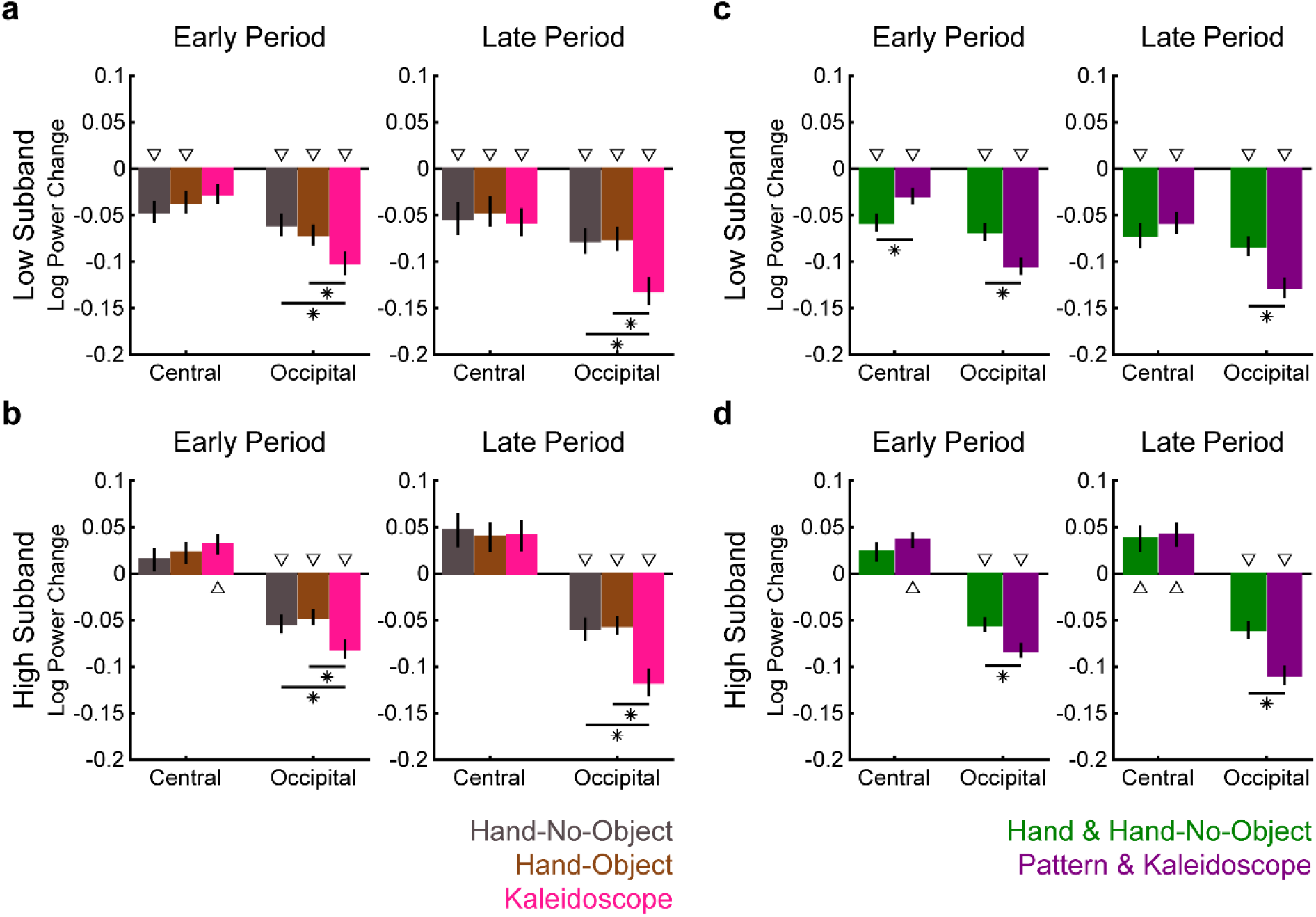
Re-analysis of the Hobson (2015) dataset and combined analyses. Mean log power change values in **(a)** Low and **(b)** High subbands in Early (left) and Late (right) periods for Hand-No-Object (HNO), Hand-Object (HO), and Kaleidoscope videos in Central and Occipital sites in the Hobson and Bishop (2016) dataset. Similarly, mean log power change values in **(c)** Low and **(d)** High subbands in Early (left) and Late (right) periods in the combined dataset, in which Hand and Pattern responses in the present study were combined with the HNO and Kaleidoscope responses, respectively, in the Hobson and Bishop (2016) dataset (n=91). Error bars depict SEMs. Asterisks indicate significant differences between pairs of stimulus categories. Triangles indicate significant differences from zero. For analyses in (a) and (b), the sample size was 61 and the significance level was Bonferroni-corrected at 0.05/6=0.008 (see Supplementary Tables 6 and 7 for all statistics). For analyses in (c) and (d), the sample size was 91 and the significance level was Bonferroni-corrected at 0.05/4=0.013 (see Supplementary Tables 8 and 9 for all statistics).

We further analyzed our findings together with the Hobson and Bishop (2016) results by combining the HNO condition with our Hand condition and the Kaleidoscope condition with our Pattern condition. The combined dataset with a total of 91 participants was analyzed via four separate 2 study (present, Hobson & Bishop) x 2 site (Central, Occipital) x 2 stimulus (Hand & HNO, Pattern & Kaleidoscope) mixed ANOVAs for Low and High subband responses in the Early and Late periods separately. Importantly, none of these ANOVAs revealed a significant interaction of study with site and/or stimulus (*F_1,89_*≤1.283, *p*≥0.260, *BF_inc_*≤2.226), and all of them indicated significant site x stimulus interactions (*F_1,89_*≥11.976, *p*≤8×10^-4^, *BF_inc_*≥107.060). In Occipital sites, in both Low and High subband responses during both the Early and Late periods, there were significant suppressions for both stimulus categories (*t_90_*≥6.357, *p*≤8×10^-9^, *BF_10_*≥10^6^; see Supplementary Table 8 for all statistics) and these suppressions were significantly stronger for Pattern and Kaleidoscope videos than Hand and HNO videos (*t_90_*≥3.910, *p*≤2×10^-4^, *BF_10_*≥111.343, Figure 6c,d; see Supplementary Table 9 for all statistics). Similarly, Central sites also yielded significant suppressions for both stimulus categories in both Early and Late periods, but only in Low subband responses (*t_90_*≥3.387, *p*≤0.001, *BF_10_*≥22.044, Figure 6c). In contrast, High subband responses in Central sites showed significant enhancements for Pattern and Kaleidoscope videos during the Early period and for both stimulus categories during the Late period (*t_90_*≥2.616, *p*≤0.010, *BF_10_*≥2.853, Figure 6d). Critically, in Central sites, there was significantly more suppression for Hand and HNO videos compared to Pattern and Kaleidoscope videos only in Low subband responses during the Early period (*t_90_*=4.062, *p*=10^-4^, *BF_10_*=183.908, Figure 6c). Taken together, the results of the two studies were not found to be different from each other and they together showed that only in the lower frequency subband, there was stronger mu suppression during observation of hand movements as compared to non-biological motion soon after the onset of the movement, while in the higher frequency subband, observation of non-biological motion can elicit mu enhancement instead of suppression.

### Fine Temporal Dynamics of Motion are Followed by Low Subband Responses for Hand, and both Low and High Subband Responses for Face Movements

We next explored to what extent Low and High subband responses in Central, Temporal, and Occipital sites were related to the motion dynamics in the Hand, Face, and Pattern videos. We performed these analyses at two different levels: within-video and between-video analyses. Within-video analyses are reported in this section and the between-video analyses are explained in the next section. Due to the repetition of the same movement in each video, our videos had motion-energy temporal profiles with two peaks as shown in Figure 1b. In addition, as mentioned above, temporal dynamics of Low subband responses depicted in Figure 3a indicated suppression profiles with two negative peaks. We investigated how well the motion and neural response temporal profiles correlated with each other at different lags. Low subband responses in Occipital sites indicated significant negative correlations between motion energy and neural activity for all stimulus categories with lags starting from 100-167 ms (Hand: 133-333 ms; Face: 167-267 ms and 600-967 ms; Pattern: 100-333 ms; Figure 7a). In contrast, there were significant negative correlations for only Hand videos in Central sites and only Face videos in Temporal sites (Figure 7a) with lags starting from 267 ms (Hand: 267-500 ms; Face: 267-800 ms). These findings suggest that mu suppression in the lower subband not only shows Hand and Face action selectivity for Central and Temporal sites, respectively, but also indicates that these suppressions follow motion dynamics in these actions with similar latencies, which together occur later than the motion-driven activity in the visual cortices. Surprisingly, at 0-100 ms lags, positive correlations were found between motion energy and Low subband responses for Face videos in Temporal sites (Figure 7a). High subband responses revealed similar results in Temporal sites (positive correlations: 0-67 ms; negative correlations: 367-733 ms; Figure 7b). In Occipital sites, there were negative correlations only for Face videos (133-267 ms and 700-733 ms), whereas, in the Central sites, there were significant positive correlations for Hand (633-700 ms) and Face (433-867 ms) videos at relatively long lags between motion dynamics and neural responses (Figure 7b).

**Figure 7.**
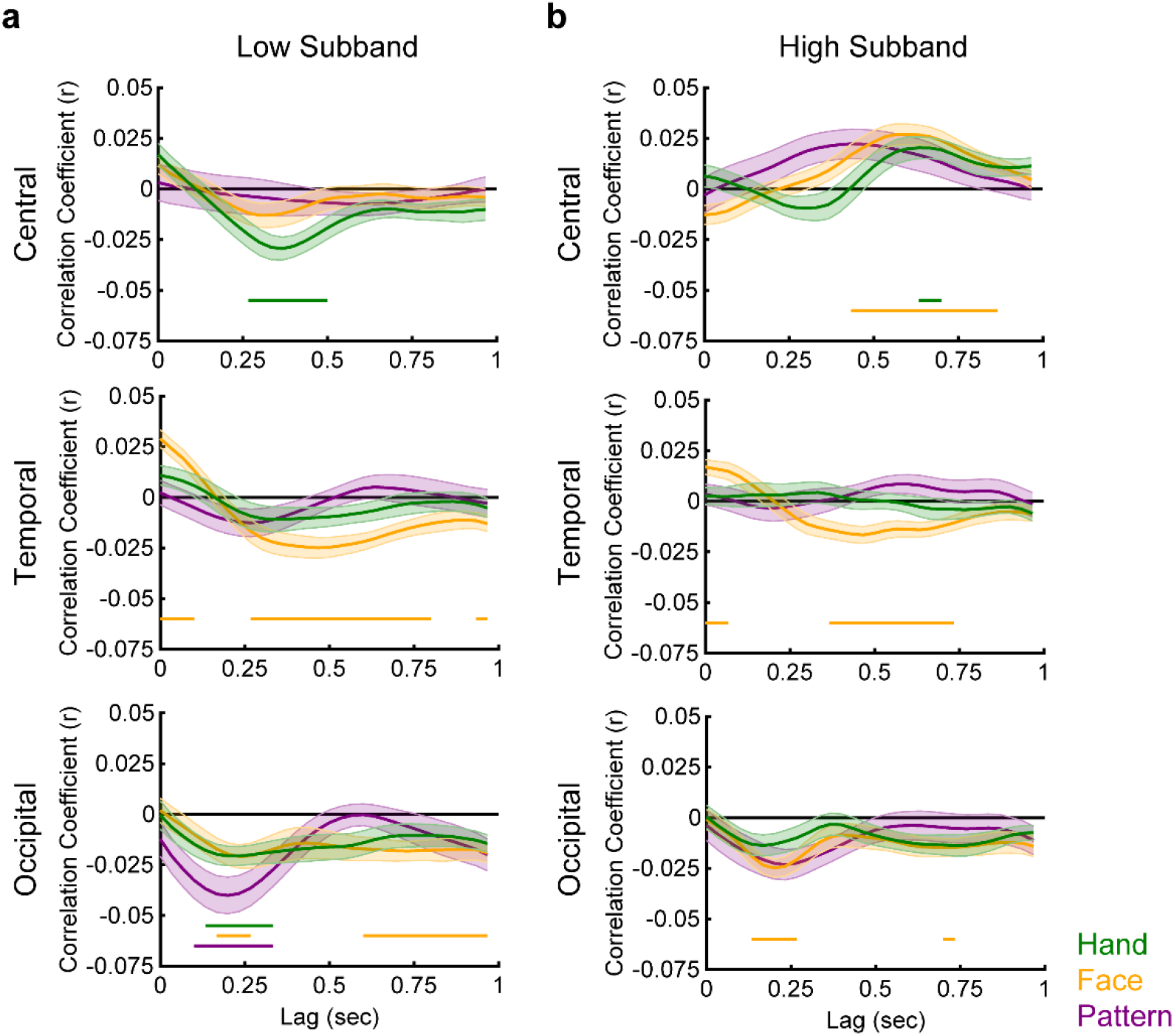
Within-video lagged correlations between neural responses and motion energy values. Mean correlation coefficients between frame-to-frame motion energy values of videos and **(a)** Low and **(b)** High subband responses at lags from 0 to 1 second in Central, Temporal, and Occipital sites. Shadings depict SEMs. Horizontal bars within individual plots indicate time windows during which Hand (green), Face (orange), and Pattern (purple) correlation coefficients were significantly lower than zero. For all analyses, the sample size was 30 and the significance level was corrected for false discovery rate at a *q* level of 0.05.

**Figure 8.**
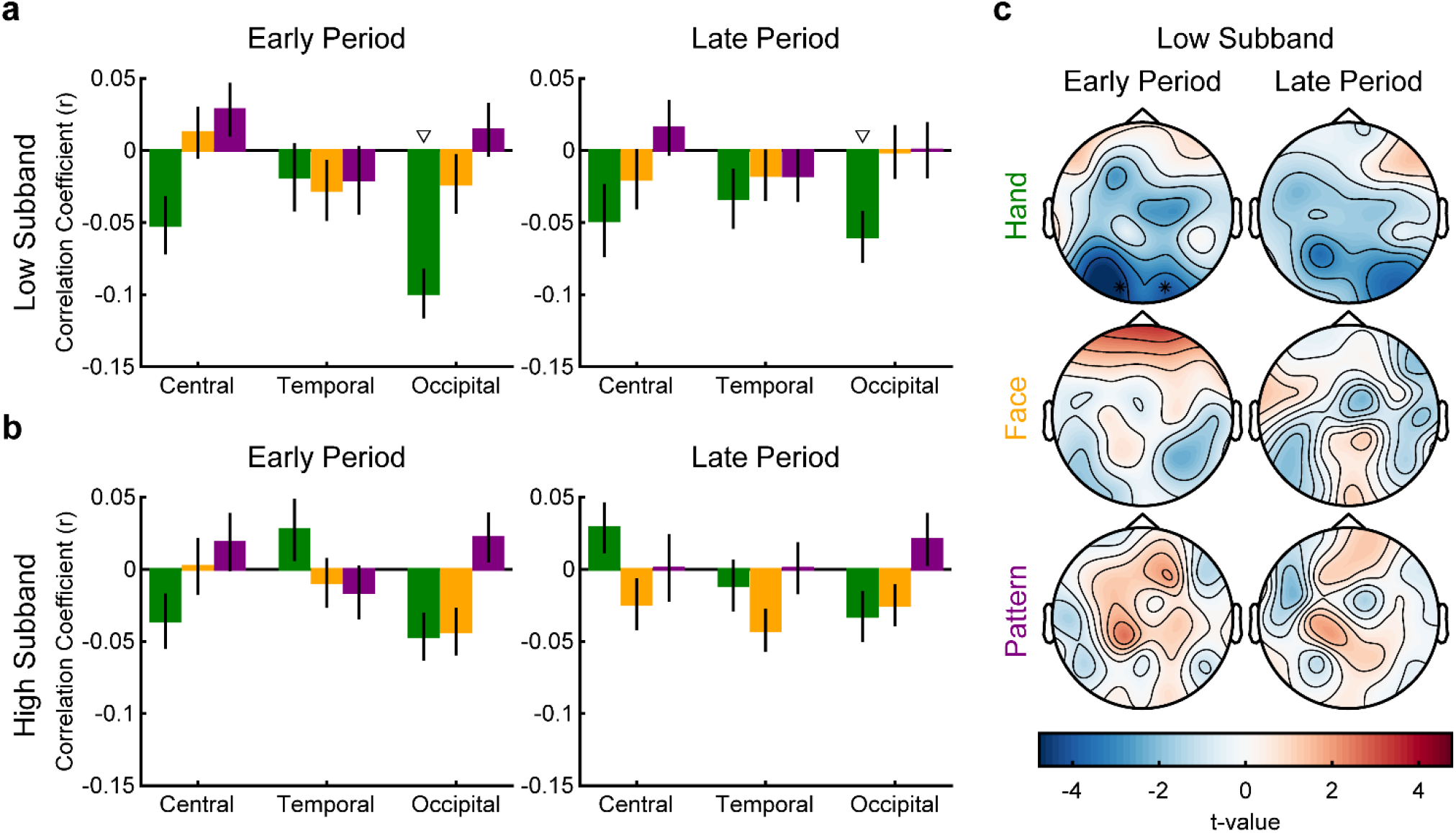
Between-video correlations between neural responses and perceived motion levels. Mean correlation coefficients between perceived motion ratings in the validation study and **(a)** Low and **(b)** High subband responses in Early (left) and Late (right) periods for Hand, Face, and Pattern videos in Central, Temporal, and Occipital sites. **(c)** T-values of one-sample t-tests comparing correlation coefficients to zero in Low subband responses in Early (left) and Late (right) periods for Hand, Face, and Pattern videos. For all analyses, the sample size was 30. For analyses in (a) and (b), triangles indicate significant differences from zero, and the significance level was Bonferroni-corrected at 0.05/9=0.006 (see Supplementary Table 10 for all statistics). A similar pattern of results was found when motion energy estimations (Supplementary Table 11) or participant’s own motion ratings (Supplementary Table 12) were used in analyses, instead of the motion ratings in the validation study. For analyses in (c), asterisks indicate significant differences from zero, and the significance level was corrected for false discovery rate at a *q* level of 0.05

### Overall Perceived Motion Levels Across Videos do not Correlate with Low or High Subband Responses

We further examined the relationship between motion and neural responses at a between-video level by calculating correlation coefficients between neural responses and the motion ratings we collected during the validation of the videos. These analyses revealed significantly negative correlations only for Hand videos in Low subband activity in Occipital sites during both Early and Late periods (*t_29_*≥3.336, *p*≤0.002, *BF_10_*≥15.705; Figure 7a,b; see Supplementary Table 10 for all statistics). Note that, although there were similar but non-significant negative correlation coefficients for Hand videos in Central sites as well, the patterns of results for different stimulus categories in Central sites were the same as, but lower than, those in Occipital sites, suggesting that the source of these effects might have been Occipital rather than Central sites. This is also reflected in the topographical distribution of correlation coefficients in Figure 7c, which shows strongly negative correlation coefficients for O1 and O2 sites and a similar but weaker trend spreading to more central regions. Similar analyses conducted separately with motion energy estimations (Supplementary Table 11) or participant’s own motion ratings (Supplementary Table 12), instead of the motion ratings in the validation study, yielded largely similar patterns of results.

### Inter-individual Variability in Low or High Subband Responses is not Explained by Interpersonal Reactivity Scores

Finally, we assessed whether the individual variability observed in neural responses can be explained by participants’ interpersonal skills as measured by the Interpersonal Reactivity Index (Davis, 1983). We first tested the degree of consistency in neural activity across different stimulus categories by calculating the correlation coefficients among pairs of stimuli. Except for a few pairs, for almost all stimulus category pairs in Low and High subband responses in Central, Temporal, and Occipital sites during Early and Late periods, there were significantly positive correlations (*r_28_*≥0.512, *p*≤0.004, *BF_10_*≥12.206; see Supplementary Table 13 for all statistics), indicating that there were inter-individual trait and/or state characteristics that affected the neural responses for all stimulus categories similarly. Nevertheless, interpersonal reactivity characteristics were not one of these critical factors that explained individual variability in neural activity, as none of the correlation coefficients between Low or High subband responses and Interpersonal Reactivity Index scores were statistically significant (*r_28_*≤0.319, *p*≥0.085, *BF_10_*≤0.932; see Supplementary Table 14 for all statistics). Similar null results were obtained when each of the four subscales of the Interpersonal Reactivity Index were entered into analyses individually.

## Discussion

While it is widely accepted that suppression of mu oscillations in central EEG sites is a homogeneous neural response within the 8-13 Hz frequency range, our data-driven approach revealed divergent findings, particularly during the observation of hand movements. Our findings clearly showed a significant suppression specifically in the lower (8-10.5 Hz), but not in the higher (10.5-13 Hz), subband in central sites. Up to date, only a handful of studies separately assessed the subbands constituting the conventional mu frequency band. Cochin et al. (1999) analyzed the subbands composing the conventional theta, alpha, and beta frequency bands and revealed a significant suppression during both the observation and execution of hand actions in central sites in the low (7.5-10.5 Hz), but not in the high (10.5-13 Hz), alpha subband. Similarly, Frenkel-Toledo et al. (2013) explored neural responses during action observation and execution of object-directed grasping movements and found more suppression in the low (8-10 Hz) than the high (10-12 Hz) alpha subband or the entire band (4-12 Hz) during action observation. Additionally, Frenkel-Toledo et al. (2014) tested stroke patients with either left or right hemisphere damage during action observation and reported that the extent of damage to the right inferior parietal lobe was significantly related to the magnitude of suppression in the low (8-10 Hz), but not in the high (10-12 Hz), subband in the C3 site, which corresponds to the unaffected hemisphere. Lastly, Dumas et al. (2014) documented a significant suppression in the low (8-10 Hz), but not in the high (11-13 Hz), subband in central sites during action observation in an autism spectrum disorder (ASD) group. In healthy participants, the C1 and C3 sites, which are typically analyzed in mu suppression studies, showed a significant suppression in the low, but not in the high, subband. Our findings corroborate these studies in specifically highlighting the low mu subband in action observation-induced suppression. During action execution, on the other hand, Frenkel-Toledo et al. (2013) reported suppression predominantly in the high alpha range (9-12 Hz). In parallel, Pfurtscheller et al. (2000) reported somatotopically selective suppression in the high subband (10-12 Hz) before and during execution of hand and foot movements and non-selective suppression in the low alpha subband. These studies suggest that suppression in the high mu subband might predominantly reflect action execution processes. In our study, we did not include an action execution condition, thus are not able to provide empirical contributions to these discussions.

Despite the low and high subband differentiation in central sites during the observation of hand actions, there was no such pattern in temporal sites during the observation of facial actions. A limited number of previous studies investigating mu suppression during the observation of facial actions all analyzed the entire 8-13 Hz frequency range without separating into subbands in central sites. Karakale et al. (2019) assessed mu suppression during the observation of static emotional and neutral facial expressions and reported significantly greater suppression compared to non-biological stimuli only for neutral expressions in central sites. Sakihara and Inagaki (2015) demonstrated mu suppression during the observation of tongue thrust movements dominantly in the left hemisphere in central regions. Although we also found significant suppression (specifically, in the low subband) in central sites in response to face actions, significantly stronger suppression in response to viewing face actions compared to non-biological movements was only observed in frontotemporal sites (see Supplementary Table 5 for the list of significant sites). These electrode sites topographically correspond to the facial region of the motor cortex and the inferior frontal gyrus, which is well-known to be involved in face perception (Kesler-West et al., 2001; Liakakis et al., 2011; Morecraft et al., 2004). Largely in line with our findings, Muthukumaraswamy et al. (2006) used magnetoencephalography (MEG) to examine suppression in oscillatory power in alpha (8-15 Hz), beta (15-35 Hz), and gamma (35-70 Hz) frequency bands during the observation of non-linguistic (e.g., biting) and linguistic (e.g., articulation of the word “seven”) orofacial actions, together with non-biological (e.g., opening and closing of a mechanical aperture) motion. The results indicated strong alpha and beta suppression, especially for object-directed non-linguistic mouth movements predominantly in lateral sensorimotor areas, which were more lateral than the sensorimotor areas that would be covered by the C1 and C3 electrodes typically used in mu suppression studies. In addition, non-linguistic mouth movements were shown to induce concurrent alpha and beta suppressions, which is conceptually in line with our findings showing no low-high frequency range differentiation for face actions. Taken together, our findings corroborate these reports by indicating selective mu suppression in more lateral than in medial sensorimotor areas during the observation of facial actions. The neural mechanisms leading to the presence and absence of clear mu subband differences in response to hand and face actions, respectively, remain unresolved. It is important to note that these oscillations, which seem homogenous across the 8-13 Hz frequency range, recorded in the frontotemporal sites cannot be the mere reflection of the alpha oscillations originating in the occipital sites given that they show stronger suppressions for Face than for Pattern stimuli, whereas the alpha oscillations in the occipital sites do not.

We also re-analyzed the dataset of Hobson and Bishop (2016), a highly cited registered report that carefully controls for attentional engagement differences across conditions and potential action-execution-related activities during action observation through electromyography (EMG) recordings. They analyzed neural responses in the 8-13 Hz frequency band and reported significant mu suppression during the observation of hand actions, but not kaleidoscope movements, in central sites using a within-trials baseline window. We prepared our stimuli and baseline-correction technique in line with their study, which enabled us to apply our analysis pipelines to their raw dataset. When we separately analyzed the mu subbands, we confirmed our findings, showing significant suppression in the low, but not in the high, subband. Furthermore, when we conducted a more robust analysis by combining their Hand-No-Object (HNO) condition with our Hand condition and their Kaleidoscope condition with our Pattern condition, we demonstrated stronger suppression for hand actions than for non-biological movements only in the low subband, whereas there were generally enhancements in the high subband. A consensus in the literature is that to be accepted as a direct indicator of mirror neuron activity, suppression in mu oscillations must be exclusively selective for the observation of internally simulatable biological actions and not seen during the observation of non-biological sources of motion (Hobson & Bishop, 2017). Nevertheless, our results showing significant suppression during the observation of non-biological motion (although generally less in magnitude than the suppression during the observation of hand actions) are in direct contradiction with this view. One possibility that can account for the present findings to preserve the conventional mirror neuron system approach, is that participants in our study were somehow able to internally simulate the non-biological pattern movements, leading to small but significant mu suppression. Although this argument is empirically impossible to rule out, another possibility, which we find more plausible, is that mu suppression is not the direct indicator of an internal simulation process.

Instead of indicating a direct action-matching mechanism, mu suppression might potentially reflect neural representations related to action possibilities that are activated during the observation of any visual input. According to this formulation, electrodes placed above the hand areas of the sensorimotor cortices show stronger low mu subband suppression during the observation of hand actions compared to any other visual input, because these inputs activate the widest array of very specific hand action possibilities that might follow the observed actions in these brain regions. On the other hand, the observation of non-biological movements, such as our pattern stimuli or kaleidoscopic movements, leads to weaker low mu subband suppression in the same electrodes, because they elicit only a limited array of general action possibilities in the same sensorimotor regions of the observers. Note that alternative explanations to not just the meaning of the mu suppression as measured by EEG, but the functioning of the mirror neuron system in general have also been provided. For instance, Csibra (2007) provided an alternative explanation to the conventional view on mirror neuron functioning - the direct matching hypothesis - which essentially suggests that we understand the goals of others’ actions primarily through automatic matching of the observed actions in the observer’s own motor system (Gallese et al., 2004; Rizzolatti et al., 2001). Csibra (2007) reviewed several lines of evidence showing that goal understanding is possible without internal simulation and argued that action understanding is done outside the motor system. Instead of directly matching what others are doing, the motor system might anticipate what others will be doing. Similar accounts have also been formulated by others (Kilner et al., 2007; Keysers & Gazzola, 2014), emphasizing the potential role of the mirror neuron system in predictive coding of what others will be doing. Nevertheless, it is highly important to emphasize that these accounts of mirror neuron system functioning can still not entirely explain our findings if mu suppression directly reflected the mirror neuron activation. It is because these accounts argue that mirror neuron activity predicts what others will be doing; for this, there still needs to be the other, whereas we found small but significant suppression even in the absence of another individual. Thus, we believe that, even if the mirror neuron system does not directly match the current actions, but predicts the future actions, of the observed individual, mu suppression cannot be a direct indicator of this functioning. One way to reconcile the two accounts would be to argue that mu suppression reflects a combination of two processes, one of which is the elicited generalized neural activity in the sensorimotor regions in response to any visual input (including non-biological motion) and the other is a direct matching-related and/or predictive coding-related neural activity selective for biological motion. The alternative explanations that we discuss about the possible mechanisms that mu suppression might reflect need to be thoroughly assessed in carefully thought experimental designs in future studies. Nevertheless, we believe that our analyses of the fine temporal profiles of mu suppression and the motion dynamics of the observed actions we discuss below might prove to be more informative than the average suppression levels in the understanding of the mechanisms that mu suppression might reflect.

We conducted the in-depth analysis of how motion dynamics of observed actions relate to mu suppression both at the within-video, fine temporal level and at the between-video, overall perceived motion level. Interestingly, these analyses revealed different results. The low alpha oscillations were tightly coupled with the fine temporal dynamics of observed actions similarly for all stimulus categories with ∼100-ms lags in occipital regions. These lags were consistent with the findings of an intracranial recording study that showed that changes in visual dynamics of movies depicting naturalistic motion could be detected within ∼100 ms in the ventral visual cortical areas (Isik et al., 2018). The lower frequency subband of the mu oscillations, on the other hand, was significantly coupled with temporal dynamics of only hand actions in central electrodes positioned above the hand region of the sensorimotor cortices and only for face actions in temporal electrodes positioned above the face region of the sensorimotor cortices with relatively longer lags around 267 ms. To the best of our knowledge, no study up to date conducted an intracranial investigation of the lags between visual motion and neural responses in sensorimotor cortical areas in humans. Mukamel et al. (2010) reported relatively longer latencies around 650 ms for the supplementary motor area single- and multi-unit responses during action observation, however, these latencies were computed as the first 100-ms time bin at which neural responses reached their maximum, not as the first time bin at which the neural responses significantly increased compared to a baseline period, and hence do not necessarily indicate the onset of neural activity during action observation. It is also highly crucial to note that the lags we report do not indicate the absolute onset times of neural firing. Instead, they represent the midpoints of the earliest time windows of lags that indicate significant relationships between neural oscillations and motion dynamics in videos. These findings show that, although mu oscillations in the low-frequency subband show specificity for different body parts in corresponding regions of the cortex, the timings of these processes in relation to motion dynamics of actions seem similar and come after the non-stimulus-specific neural responses in occipital regions. Whether these timing differences indicate a process by which the early motion-dependent activity in the occipital regions is gated and supplied to specific parts of the sensorimotor cortical areas for specific body parts or the body part-selective activity in the sensorimotor areas emerge independently of the early occipital activity remains to be resolved by studies utilizing simultaneous neuromodulation and measurement techniques.

The results of our between-video analyses were not in line with those of the within-video analyses. There was no significant relationship between the average perceived motion levels depicted in the videos and the average magnitudes of mu suppression except for the low subband responses to Hand videos in the occipital sites. Note that the weak but similar patterns of findings in central sites are likely to be mere reflections of the activity in the occipital sites rather than reflecting local processing in the sensorimotor regions. The observed results remained consistent even when the correlation analyses were conducted with objectively calculated average motion levels or participants’ own perceived motion ratings. Taken together, our within- and between-video analyses clearly indicate that the low mu subband activity reflects motion dynamics in observed actions relatively within a local action sequence, but not the overall motion levels absolutely across the observation of different action sequences.

Finally, we found that neural responses to different stimuli were strongly correlated with each other across participants, indicating trait and/or state characteristics that influence the processing of various sources of motion similarly. Nevertheless, neither the generalized empathic skills as quantified by the total Interpersonal Reactivity Index (IRI) scores, nor any of its empathic concern, fantasy, personal distress, or perspective-taking subcomponents were related to these individual differences in neural responses. Other studies revealed mixed results regarding the relationship between mu suppression and empathic skills as measured by the IRI, some reporting significant relationships with personal distress (Cheng et al., 2008; Yang et al., 2009; Martin et al., 2017; Peled-Avron et al., 2016), empathic concern (Woodruff & Klein, 2013; Brown et al., 2016) and fantasy subscales (Neufeld et al., 2016), while others showing no relationship (Woodruff et al., 2011; Perry et al., 2010; Silas et al., 2010; Lübke et al., 2020; Milston et al., 2013; McCormick et al., 2012; Horan et al., 2014). All these studies used fairly different stimuli, designs, and analyses; however, taken together our findings are in line with the ladder group of studies. Nevertheless, it is important to note that, in a high-powered study (n=252), DiGirolamo et al. (2019) showed that, although there was no significant relationship between mu suppression and any of the IRI subcomponents at the electrode level, when analyzed at the component level, greater mu suppression was associated with greater empathic concern scores in the IRI. We analyzed our dataset only at the electrode level and thus are agnostic to the possible effects that would be revealed at the component level. Future studies will need to assess these effects systematically to arrive at conclusive results as to whether part of the inter-individual variability in mu suppression can be explained by variations in empathic abilities.

## Materials and Methods

### Participants

A total of 42 participants with normal or corrected vision and without any current diagnosis of neurological or somatic disorders were recruited to participate in the present study. Two participants were excluded due to technical problems during EEG recordings. Another 10 participants were excluded from the EEG analyses due to not performing successfully in the motion rating task (see *Results* for details). Thus, a final sample of 30 participants aged between 19 and 34 years old (*Mean*=22, *SD*=3; female/male=24/6) was included in all statistical analyses. All procedures were approved by the Kadir Has University Human Research Ethics Committee and each participant read and signed a written informed consent according to the Declaration of Helsinki before participating in the study.

### Stimuli

A novel stimulus set consisting of 156 grayscale videos depicting hand, face, or pattern movements was generated to be utilized in the present study. Hand videos displayed 12 different simple finger movements (e.g., index finger performing a lateral movement, all fingers opening and closing; see Supplementary Table 1 for the full list of movements), together with a no-movement condition, seen from the dorsal view of the right hand, each performed by a female and a male actor and recorded on a white background (Figure 1a). Similarly, Face videos consisted of 12 different facial movements (e.g., eyes blinking, jaw performing a chewing action; see Supplementary Table 1 for the full list of movements) and a no-movement condition, performed by a female and a male actor (Figure 1a). Each movement in the Hand and Face videos was repeated twice at a once-per-second rate, the performance of which was aided by a metronome during the video recordings. Two-second videos were cut, starting from the last frame before the onset of the first epoch of the movements to ensure that the baseline views of hands and faces were in their resting positions. Videos in which the hand or the face in the last frame in the two-second versions of the videos were not back in their resting positions were discarded, so that the ending of movements were also standardized. All videos were cropped, centered, and resampled such that the height of the hands (estimated from the tip of the fingers to the wrist) and faces (estimated from the tip of the hairline to the tip of the chin) occupied the center 540 pixels of the 1080×1080-pixel final videos.

To compare the hand and face movements to non-biological motion, a novel control stimulus set of different pattern movements was generated using custom computer algorithms. Clouds of either circles or squares (to create two different versions parallel to the Female and Male versions in Hand and Face videos) of different sizes and shadings were generated in a circular distribution that spread within the center ∼540×540 pixels of the 1080×1080-pixel videos (Figure 1a). A large pool of videos, in which different proportions of randomly determined individual units made linear movements of various distances, were generated. To mimic hand and face movements, pattern movements were made in a back-and-forth manner at a once-per-second rate. From the resulting large pool of 2-second videos, 13 circles and 13 squares videos were selected based on the average amount of motion energy they contained. Motion energy was calculated by taking the difference of the grayscale values of each corresponding pixel in consecutive frames of the video and then averaging across all pixels and frames. Note that only pixels within a circle with a diameter of 540 pixels centered in the middle of the videos were taken into account during the calculation of motion energy since the inclusion of background pixels with no movements would have led to an underestimation of motion. Although there are other algorithms for quantifying hand and face movements, we chose the above-mentioned model-free method of estimating motion energy to suit our need for utilizing it similarly for both biological and non-biological forms of motion. The final set of Pattern videos was selected such that the average motion energies across the videos were not different than those of the Hand and Face videos (*F_2,75_*=0.754, *p*=0.474, *BF_inc_*= 0.199; Figure 1b). It is important to emphasize that, with these motion energy estimations, we do not argue that we equated how much motion the Hand, Face, and Pattern videos had; we simply used a common metric to represent motion level variabilities within each of the three video stimulus sets.

Secondary versions of all videos were created by horizontally flipping them to control for potential lateral differences. Finally, the first frames of the videos were repeated for 4 seconds to create 4-second baselines and the last frames were repeated for 2 seconds to create 2-second post-stimulus periods, in line with Hobson and Bishop (2016). Thus, the final stimulus set consisted of 3 stimulus (Hand, Face, Pattern) x 13 movement x 2 type (Female and Male for Hand and Face videos, Circles and Squares for Pattern videos) x 2 version (Original, Horizontally flipped) = 156 videos consisting of 4-second static baseline + 2-second dynamic movement + 2-second static post-stimulus period = 8 seconds.

### Procedure

#### Motion Rating Task

The motion rating task used during EEG recordings was programmed in PsychoPy (version 2022.2.4; Peirce et al., 2019). The task consisted of two repetitions of each of the 156 video stimuli for a total of 312 trials presented in four blocks. Each trial of the task started with the presentation of a plus symbol in the center of the screen for fixation for 1 second, followed by the presentation of an 8-second video pseudo-randomly drawn from the full stimulus set such that the same stimulus, type, or version would not appear in more than four consecutive trials and the same movement would not appear in more than three consecutive trials. In 78 trials (1/4 of all trials), after the offset of the video, a motion rating screen was shown. This screen asked participants to answer the question “How much motion did the video contain?” by reporting their ratings on a 9-point scale from 1 (very little) through 5 (medium) to 9 (very much). Participants reported their ratings at their own pace by using the arrow keys on the keyboard with their right index and middle fingers to minimize movement. The rating trials were pseudo-randomly selected such that one of the four presentations (original and the horizontally flipped version, each presented twice) of each unique video was rated once. In addition, there were no more than three rating trials in a row and there was at least one rating trial in every seven consecutive trial blocks from the beginning of the task. In the remaining 234 trials (3/4 of all trials) the offset of the video was followed by the intertrial interval showing a blank black screen for 0.5 seconds. A monitor with a 47.6×26.8-cm physical size and 1920×1080-pixels resolution was used. Participants were seated ∼115 cm away from the monitor, which yielded ∼6.7 degrees of visual angle for the center 540×540 pixels of the videos that contained the majority of the motion in the videos. At the beginning of the task, the participants were instructed that they would be shown videos of hand, face, and pattern motions and would be asked to rate how much motion these videos contained in some of the trials. There was no practice trial. We chose to collect motion ratings only in a quarter of all trials to not make the entire EEG recording session very long and to not have potential movement artifacts due to rating behavior in the majority of the trials. There were no cues as to which video would be eventually rated for motion until the end of the video presentation, thus participants had to attend all videos similarly.

An alternative version of the motion rating task was used in a stimulus validation study prior to the EEG study. The stimuli and the task dynamics were all the same except the following: A blocked design was used for each stimulus category and the block order was counterbalanced across participants. Only the 2-second active period of each video was shown to participants. Each video presentation was followed by a rating screen. A 1-second intertrial interval was used. A monitor with a 52.7×29.7-cm physical size was used. The participants were seated ∼60 cm away from the monitor, which yielded ∼14 degrees of visual angle for the center 540×540 pixels of the videos. The mouse was used to collect motion ratings. Finally, before the actual ratings in each block, there was a practice phase consisting of seven videos from the stimulus set.

#### EEG Recordings

EEG recordings were collected in a Faraday cage using the Brain Vision actiCHamp system (Brain Products, Munich, Germany) from 32 active electrodes placed in a cap following the international 10-20 system. TP10 electrode was used as the reference electrode during the recordings. Before the recordings, impedances of all electrodes were brought to less than 10 kΩ by applying sufficient amounts of electrolytic gel and were verified again after the end of the recordings. Continuous EEG recordings were obtained at a 1000-Hz sampling rate with no online filters while the participants completed the motion rating task. Video onsets were detected via white pixels positioned in the bottom right corner of the videos, which were captured by a photosensor attached to the monitor. These white pixels were hidden by the photosensor and thus were completely invisible to the participants. EEG recordings lasted ∼1 hour on average.

#### Interpersonal Reactivity Index

After the EEG recordings, participants completed the Turkish version of the Interpersonal Reactivity Index (Engeler & Yargıç, 2007) originally developed by Davis (1983) for a multi-dimensional assessment of empathy. It is a widely used self-report questionnaire consisting of a total of 28 items divided into four subscales: Empathic Concern, Fantasy, Personal Distress, and Perspective-taking. Participants rated each item on a 5-point scale from 0 (does not describe me well) to 4 (describes me very well). The average of all 28 items was used as the interpersonal reactivity score of each participant. Participants’ scores for each of the four subscales were also estimated separately.

### Data Analysis

#### General Statistical Approach

Data preprocessing and scientific computations were conducted using the FieldTrip toolbox (version 20230402; Oostenveld et al., 2011) and custom scripts in MATLAB (version R2022B), and statistical analyses were conducted in JASP (version 0.16.4; JASP Team, 2022). The details of the statistical tests are described below and in the *Results* section specifically for each analysis. Alpha level was set to 0.05 but specifically corrected via permutation-based cluster-correction for analyses on time-frequency profiles (Maris & Oostenveld, 2007), via false discovery rate-based correction for analyses with >30 temporally or spatially dependent statistical tests (Benjamini & Yekutieli, 2001), and Bonferroni correction for analyses with <10 planned comparisons. The details of each of these correction methods are described specifically for each analysis below. Greenhouse–Geisser-corrected *p* values were used for evaluation of statistical significance when sphericity assumptions were violated as indicated by significant Mauchly’s tests of sphericity. We supplemented the frequentist analyses with Bayesian statistical analyses when possible. *BF_10_* represents the Bayes Factor indicating the ratio of the evidence in support of H_1_ to the evidence for H_0_ in paired-samples and one-sample Bayesian t-tests. *BF_inc_* represents the Bayes Factor indicating the ratio of the evidence when a factor is included compared to the evidence when it is excluded from the matched models in repeated-measures Bayesian ANOVAs. *BF* values greater than 3 indicate significant evidence for there being an effect or difference, as in *p*<0.05, whereas *BF* values smaller than 1/3 provide significant evidence for there being no effect or difference, that is, evidence for the absence of an effect or difference (Keysers et al., 2020). This property of Bayesian statistical analysis makes it superior to frequentist statistical tests, in which a *p* value greater than the alpha level cannot be used to conclude whether there is evidence for the absence of an effect (support for H_0_) or absence of the evidence for an effect (inconclusive test).

#### Motion Ratings

For each participant, the correlation between that participant’s own motion ratings and the average motion ratings in the validation study across all videos was calculated via a two-tailed non-parametric Kendall’s tau correlation coefficient analysis. Nonparametric tests were preferred due to the ordinal nature of the ratings. The alpha level of 0.05 was not corrected for multiple comparisons for these analyses. Ten participants who failed to show significant correlations were excluded from all further analyses. Similar correlation coefficients were also calculated for Hand, Face, and Pattern videos separately (see Supplementary Table 15 for the full list of correlation coefficients). These correlation coefficients were compared among the stimulus categories in a 3 stimulus (Hand, Face, Pattern) repeated-measures ANOVA. In addition to these correlation coefficients, the raw motion ratings were also compared similarly in two separate 3 stimulus (Hand, Face, Pattern) repeated-measures ANOVAs, one for the EEG study and one for the validation study. Significant effects in these analyses were further probed via 3 separate planned comparisons for all possible stimulus category pairs via two-tailed paired-samples t-tests. For each of these 3 tests, a Bonferroni-corrected alpha level of 0.05/3=0.0167 was used.

#### EEG Preprocessing

Raw EEG data was high-pass filtered at 0.1 Hz and segmented into trials of 8 seconds from the onset to the offset of the videos. Eye blink artifacts were first detected automatically at a relatively sensitive z-score of 3 from the Fp1 and Fp2 electrodes, which yielded high levels of false positives. Thus, these detections were then manually accepted or rejected by a trained operator blind to the trial identities. Muscle artifacts were also detected by the same operator. Trials were rejected if any of the detected artifacts fell within the 2-6 seconds of the videos since this time window was used in all analyses. Across participants, 2% to 70% of all trials were rejected during this stage of preprocessing (*Mean*=33%, *SD*=21%; see Supplementary Table 15 for the full list of number of trials included in statistical analyses). Finally, voltage values were converted to current source density (CSD) values using the CSD Toolbox (version 1.1; Kayser & Tenke, 2006a, 2006b) in MATLAB. These values minimize volume-based problems in EEG and provide a reference-free representation of EEG activity.

#### Time-Frequency Decomposition

The power of CSD values in each trial was estimated from 0.25 to 30 Hz (in steps of 0.25 Hz) and from 0 to 7.990 seconds (in steps of 10 ms) using a sliding Hanning taper with a duration equal to eight full cycles for each frequency (minimum=8 seconds). Power values were then converted to logarithms in base 10. Time-frequency points that were ± 5 standard deviations from the mean of the same time-frequency points in all trials were excluded from further analyses, which was very rare (*Mean*=0.046%, *SD*=0.012%). Then, for each frequency, all log power values were subtracted from the average log power of the baseline period, which was the 2-4 seconds of the videos. Finally, for each participant, the average of the time-frequency representations for Hand, Face, and Pattern videos were calculated separately. Statistical analyses on time-frequency representations were performed on these average log power change values.

#### Statistical Analyses on Time-Frequency Representations of Responses

We performed two related, but separate, sets of statistical analyses on time-frequency representations. The first set of analyses aimed at answering whether there were time-frequency points that showed significant suppression or enhancement of log power change values. These analyses were conducted separately for Hand, Face, and Pattern videos. In the first stage of these analyses, for each time-frequency point separately, a two-tailed one-sample t-test was used to compare the log power change values of all 30 participants to zero at an alpha value of 0.05. The statistical significance of the clusters consisting of individually significant time-frequency points was then assessed using a permutation-based cluster-correction method (Maris & Oostenveld, 2007). For each analysis, we performed 1000 iterations, each of which was used to estimate the maximum cluster sum that would be expected under the null hypothesis. Then, the resampling distribution consisting of these 1000 values was used to calculate the probability of each of the cluster sums observed in the actual dataset under the null hypothesis. A two-tailed alpha value of 0.05 was used for assessing the statistical significance of these clusters.

The second set of analyses employed the same permutation-based cluster-correction strategy but focused on pairwise stimulus comparisons. That is, in the first stage of analyses, for each time-frequency point separately, a two-tailed paired-samples t-test was used to compare the log power change values of all 30 participants between Hand and Pattern, Face and Pattern, and Hand and Face stimulus comparisons, separately. The rest of the cluster-correction method was performed as described above.

#### Statistical Analyses on Temporal Profiles of Frequency-Averaged Responses

Time-frequency representation analyses indicated marked differences between the Low (8-10.5 Hz) and High (10.5-13 Hz) subband responses, especially in Central sites. Thus, log power change values within each of these frequency bands were averaged across their corresponding frequencies, which yielded the temporal profiles of log power change values in Low and High subbands. For each of the two frequency bands and the three recording sites separately, time points in which Hand or Face responses were significantly different than Pattern responses were investigated via individual two-tailed paired-samples t-tests comparing the log power change values of all 30 participants between Hand and Pattern and Face and Pattern responses across 400 time points (4 seconds at a sampling rate of 100 Hz). For each analysis, the statistical significance of these 400 individual tests was corrected for false discovery rate at a *q* value of 0.05 and with the method that is guaranteed to be accurate for any test dependency structure (Benjamini & Yekutieli, 2001).

#### Statistical Analyses on Time- and Frequency-Averaged Responses

Temporal profile analyses indicated some differences between the Early (4-5 seconds) and Late (5-6 seconds) periods. Thus, the log power change values of all 30 participants in Low and High subbands during each of these time periods were averaged across their corresponding time points, which yielded the time- and frequency-averaged log power change values. The entire dataset was first analyzed via a 2 subband (Low, High) x 2 period (Early, Late) x 3 site (Central, Temporal, Occipital) x 3 stimulus (Hand, Face, Pattern) repeated-measures ANOVA. Factors subband and period were found to be interacting with several other factors. Thus, the two subbands and the two periods were further investigated via four separate 3 site (Central, Temporal, Occipital) x 3 stimulus (Hand, Face, Pattern) repeated-measures ANOVAs. For each of these ANOVAs, a significant interaction was further investigated via 9 separate two-tailed paired-samples t-tests for planned comparisons, contrasting the three pairwise stimulus comparisons in each of the three recording sites. For each of these 9 tests, a Bonferroni-corrected alpha level of 0.05/9=0.0056 was used. In addition to these planned stimulus comparisons, the log power change values for each of the three stimulus categories in each of the three recording sites were separately compared to zero via two-tailed one-sample t-tests to assess whether there was significant suppression or enhancement of neural responses. Similarly, for each of these 9 tests, a Bonferroni-corrected alpha level of 0.0056 was used.

The time- and frequency-averaged log power change values were also analyzed at all 32 recording sites to investigate the topographical distribution of the observed effects. The first set of these analyses compared the log power change values of all 30 participants to zero via separate two-tailed one-sample t-tests. For each stimulus category in each period and each subband, 32 tests were performed, one for each recording site. The statistical significance of these 32 individual tests was corrected for false discovery rate at a *q* value of 0.05 and with the method that is guaranteed to be accurate for any test dependency structure (Benjamini & Yekutieli, 2001). The second set of these analyses employed the same multiple testing and correction strategy but was performed with two-tailed paired-samples t-tests comparing the log power change values of all 30 participants between Hand and Pattern, Face and Pattern, and Hand and Face trials, separately.

#### Re-analysis of the Hobson (2015) Dataset

The raw data files, as well as the analysis scripts, of the study reported by Hobson and Bishop (2016) were shared in public domains (Hobson, 2015; https://osf.io/yajkz/). Raw EEG files were preprocessed using the shared scripts in the same way as described in Hobson and Bishop (2016) until the end of the conversion of the raw voltage values to CSD values. Then, these CSD values were processed with our time-frequency decomposition and time- and frequency-averaged log power change value calculation routines as described above. The entire dataset of 61 participants was first analyzed via a 2 subband (Low, High) x 2 period (Early, Late) x 2 site (Central, Occipital) x 3 stimulus (Hand-No-Object [HNO], Hand-Object [HO], Kaleidoscope) repeated-measures ANOVA. Factors subband and period were found to be interacting with several other factors. Thus, the two subbands and the two periods were further investigated via four separate 2 site (Central, Occipital) x 3 stimulus (HNO, HO, Kaleidoscope) repeated-measures ANOVAs. For each of these ANOVAs, a significant interaction was further investigated via 6 separate two-tailed paired-samples t-tests for planned comparisons, contrasting the three pairwise stimulus comparisons in each of the two recording sites. For each of these 6 tests, a Bonferroni-corrected alpha level of 0.05/6=0.0083 was used. In addition to these planned stimulus comparisons, the log power change values for each of the three stimulus categories in each of the two recording sites were separately compared to zero via two-tailed one-sample t-tests. Similarly, for each of these 6 tests, a Bonferroni-corrected alpha level of 0.0083 was used. Note that Temporal sites were not included in these analyses, because there were no Face videos that would be expected to yield selective suppression in the T7 and T8 electrodes.

Parts of the present and the Hobson (2015) datasets were also analyzed together by combining the HNO condition with our Hand condition and the Kaleidoscope condition with our Pattern condition. The combined dataset of all 91 participants was analyzed via four separate 2 study (present, Hobson & Bishop) x 2 site (Central, Occipital) x 2 stimulus (Hand & HNO, Pattern & Kaleidoscope) mixed ANOVAs for Low and High subband responses in Early and Late periods. For each of these ANOVAs, a significant interaction between site and stimulus was further investigated via 2 separate two-tailed paired-samples t-tests for planned comparisons, contrasting the two stimulus categories in each of the two recording sites. For each of these 2 tests, a Bonferroni-corrected alpha level of 0.05/2=0.025 was used. The log power change values for each of the two stimulus categories in each of the two recording sites were separately compared to zero via two-tailed one-sample t-tests. For each of these 4 tests, a Bonferroni-corrected alpha level of 0.05/4=0.0125 was used.

#### Within-video Analysis of Correlations Between Motion and Neural Responses

Lagged correlation coefficients between the temporal profiles of neural responses and the motion energy values of videos were calculated within each trial separately as follows. First, the temporal profiles of neural responses were downsampled from their original sampling rate of 100 Hz to 30 Hz, which was the sampling rate of the videos. Then, within each trial, the temporal profile of the motion energy values during the 2-second active part of the videos was taken into 30 separate Pearson’s correlation analyses, one for each of the 30 lags from 0 to 0.97 seconds in steps of 0.033 seconds, which corresponds to the steps of one frame in a sampling rate of 30 Hz. At the 0-second lag, the temporal profile of the 2-second neural response starting at the onset of the active part of the video was taken into the correlation analyses. At the 0.033-second lag, the temporal profile of the 2-second neural response starting 0.033 seconds after the onset of the active part of the video was taken into the analyses, and so on. These correlation coefficients were then averaged across the trials of the same stimulus category for Hand, Face, and Pattern videos separately. These correlation analyses were performed for Low and High subband responses in Central, Temporal, and Occipital sites, separately. To assess which lags yielded significant correlations across the participants, the correlation coefficient values of all 30 participants were compared to zero via 30 separate two-tailed one-sample t-tests, one for each lag. For each analysis, the statistical significance of these 30 individual tests was corrected for false discovery rate at a *q* value of 0.05 and with the method that is guaranteed to be accurate for any test dependency structure (Benjamini & Yekutieli, 2001). Note that significant negative correlations indicate that greater motion in videos is associated with stronger suppression in neural activity.

#### Between-video Analysis of Correlations Between Motion and Neural Responses

The relationship between motion dynamics and the neural responses was also investigated at the between-video level. For each participant, the relationship between the perceived motion ratings obtained from the validation study and that participant’s time- and frequency-averaged log power change values were quantified via Pearson correlation coefficients. These correlations were estimated for Hand, Face, and Pattern videos in Central, Temporal, and Occipital sites for Low and High subband responses in the Early and Late periods, separately. For each analysis, the correlation coefficients were compared to zero via two-tailed one-sample t-tests. For each of the sets of 9 tests, a Bonferroni-corrected alpha level of 0.05/9=0.0056 was used. The same analyses were also performed by taking into calculation the motion energy estimations or participant’s own motion ratings, instead of the motion ratings in the validation study. In the analysis with motion energy estimations, time-averaged motion energy values of videos were used in Pearson correlation coefficient estimations. In the analysis with the participant’s own motion ratings, the participant’s ratings given to each unique video were also used for the horizontally flipped version of the same video and the other repetitions of each of these videos that were not explicitly rated by the participant. This way, although participants rated only a quarter of all trials, all trials were included in Pearson correlation coefficient estimations.

#### Correlation Analyses Between Neural Responses and Interpersonal Reactivity Scores

To assess the similarity of responses to different stimulus categories, Pearson correlation coefficients were calculated between the Hand-Pattern, Face-Pattern, and Hand-Face stimulus pairs in Central, Temporal, and Occipital sites for Low and High subband responses in the Early and the Late periods separately. These correlation coefficients were compared to zero via two-tailed one-sample t-tests. For each of these 9 tests, a Bonferroni-corrected alpha level of 0.05/9=0.0056 was used. Finally, the relationship between the total Interpersonal Reactivity Index scores and neural responses was analyzed via Pearson correlation coefficients for Hand, Face, and Pattern videos in Central, Temporal, and Occipital sites for Low and High subband responses in the Early and Late periods separately. These correlation coefficients were also compared to zero via two-tailed one-sample t-tests, each of which used a Bonferroni-corrected alpha level of 0.0056. Similar analyses were also conducted on four of the subscales of the Interpersonal Reactivity Index separately.

## Author Contributions

Conceptualization: ES

Data Curation: ANB, ES

Formal Analysis: ANB, ES

Funding Acquisition: ES

Investigation: ANB

Methodology: ANB, ES

Project Administration: ES

Resources: ANB, ES

Software: ES

Supervision: ES

Validation: ES

Visualization: ANB, ES

Writing – Original Draft and Preparation: ANB, ES

Writing – Review and Editing: ANB, ES

## Acknowledgments

We would like to thank Pınar Demir and Berfin Mısırlı for their assistance in data collection and Barış Kaan Ok, Çağla Nur Demirkan, Elif Cantürk, Nurdem Okur, and Yaren Beyza Türkmen for their assistance in the preparation of the stimulus set.

## Data Availability Statement

The data that support the findings of this study are openly available at Open Science Framework (https://doi.org/10.17605/OSF.IO/QA4FR).

