## Supplementary Files for "Mu Suppression During Action Observation Only in the Lower, not in the Higher, Frequency Subband"

### Supplementary Figure 1

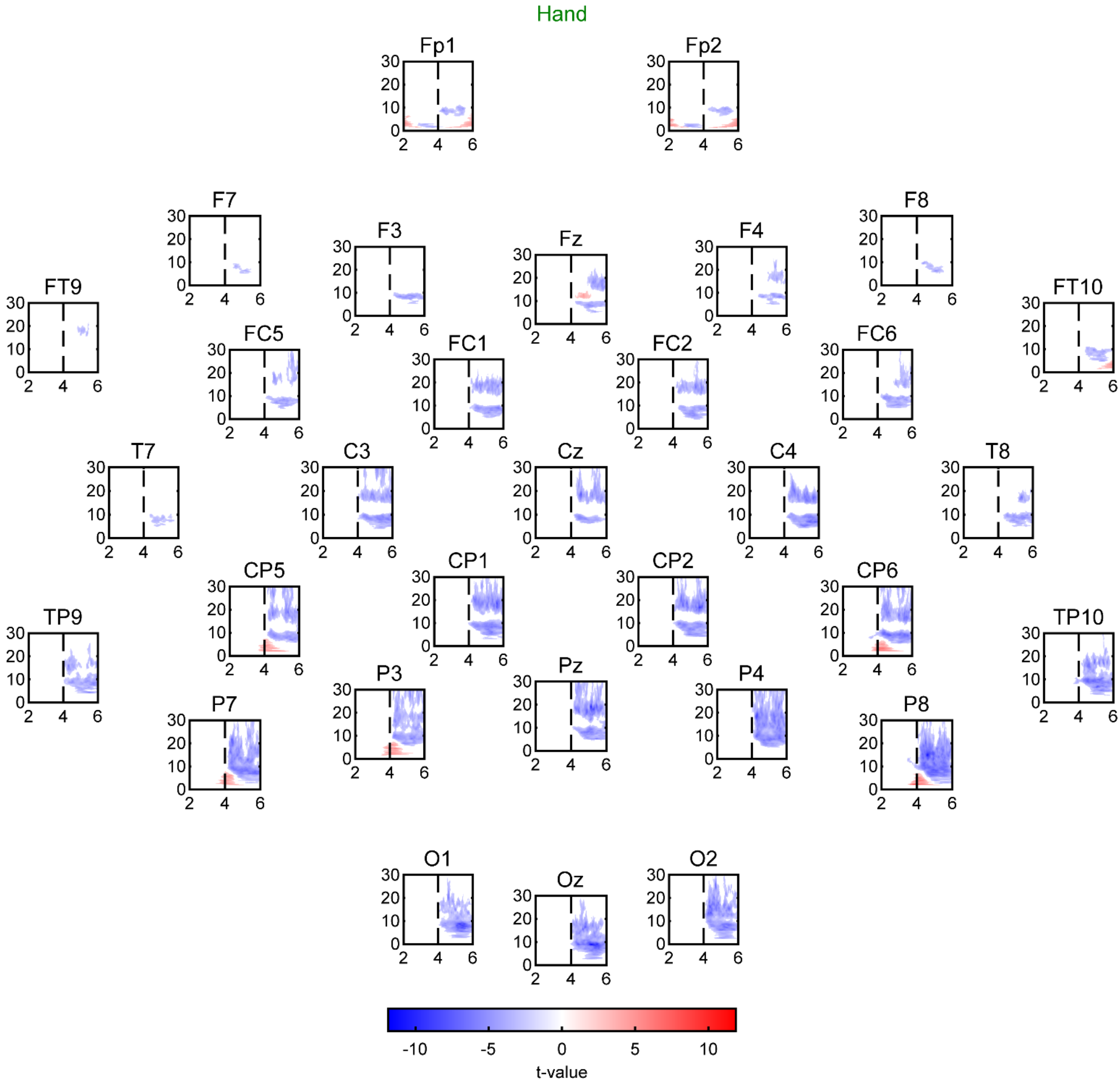


**Supplementary Figure 1. Time-frequency representations of Hand responses in 32 recording sites.** T-values of one-sample t-tests comparing log power change values to zero across frequencies and time points for Hand videos in 32 recording sites. For all analyses, the sample size was 30 and the statistical significance was determined via cluster-correction. T-values of only significant clusters are shown. Note that, during the initial 2 seconds, the video was static, and during the ladder 2 seconds, the video displayed motion.

### Supplementary Figure 2

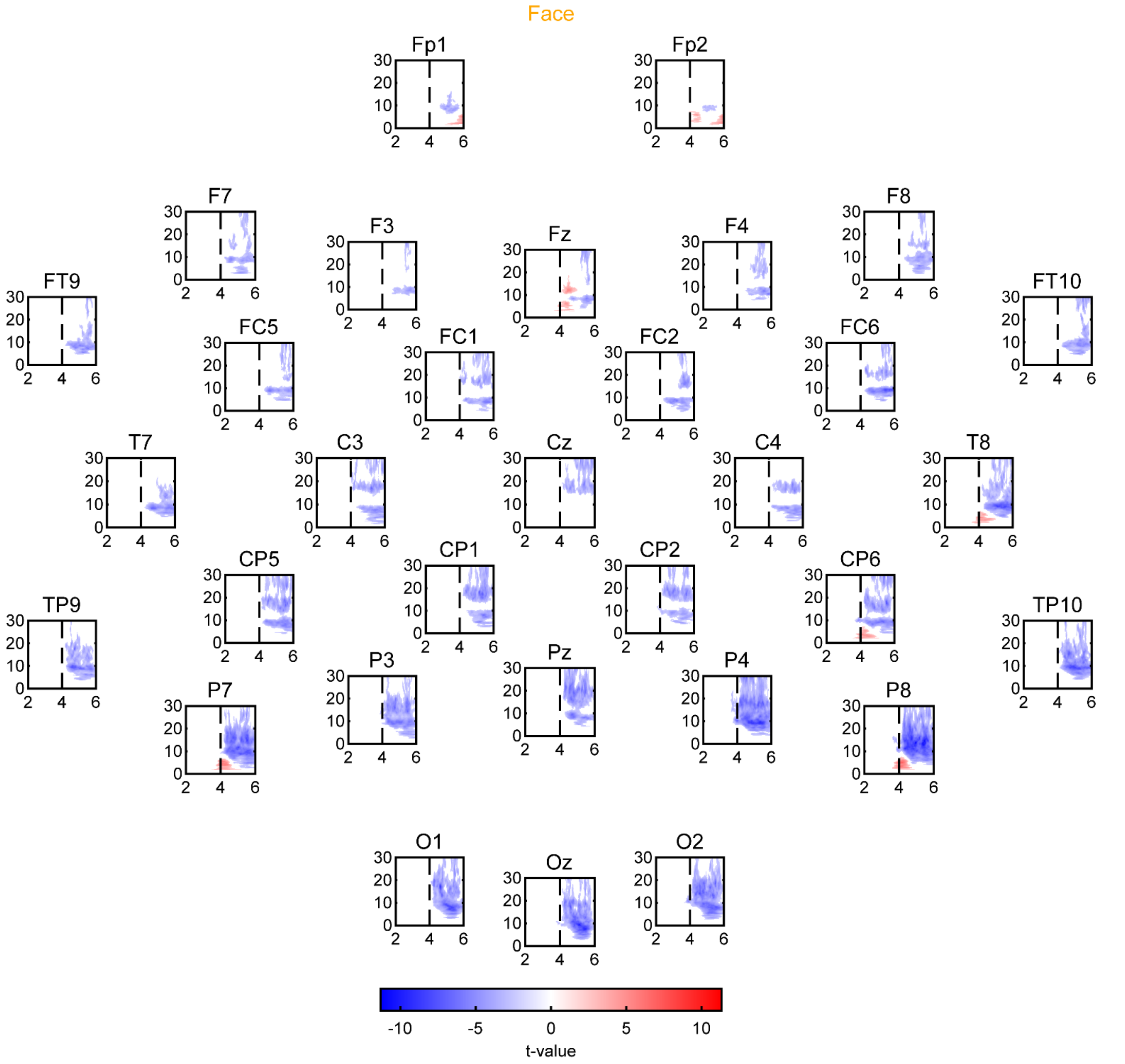


**Supplementary Figure 2. Time-frequency representations of Face responses in 32 recording sites.** T-values of one-sample t-tests comparing log power change values to zero across frequencies and time points for Face videos in 32 recording sites. For all analyses, the sample size was 30 and the statistical significance was determined via cluster-correction. T-values of only significant clusters are shown. Note that, during the initial 2 seconds, the video was static, and during the ladder 2 seconds, the video displayed motion.

### Supplementary Figure 3

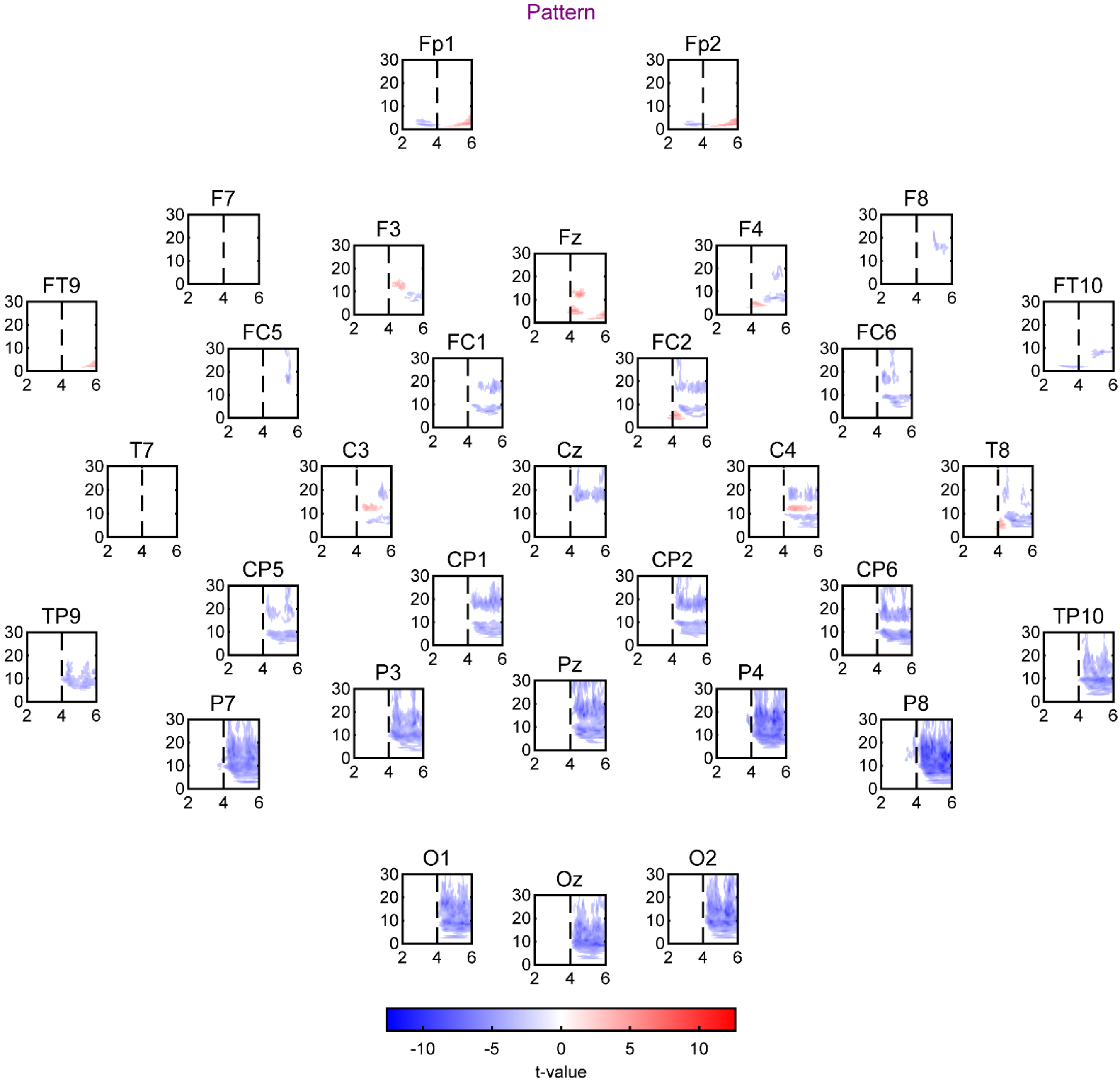


**Supplementary Figure 3. Time-frequency representations of Pattern responses in 32 recording sites.** T-values of one-sample t-tests comparing log power change values to zero across frequencies and time points for Pattern videos in 32 recording sites. For all analyses, the sample size was 30 and the statistical significance was determined via cluster correction. T-values of only significant clusters are shown. Note that, during the initial 2 seconds, the video was static, and during the ladder 2 seconds, the video displayed motion.

**Supplementary Figure 4**

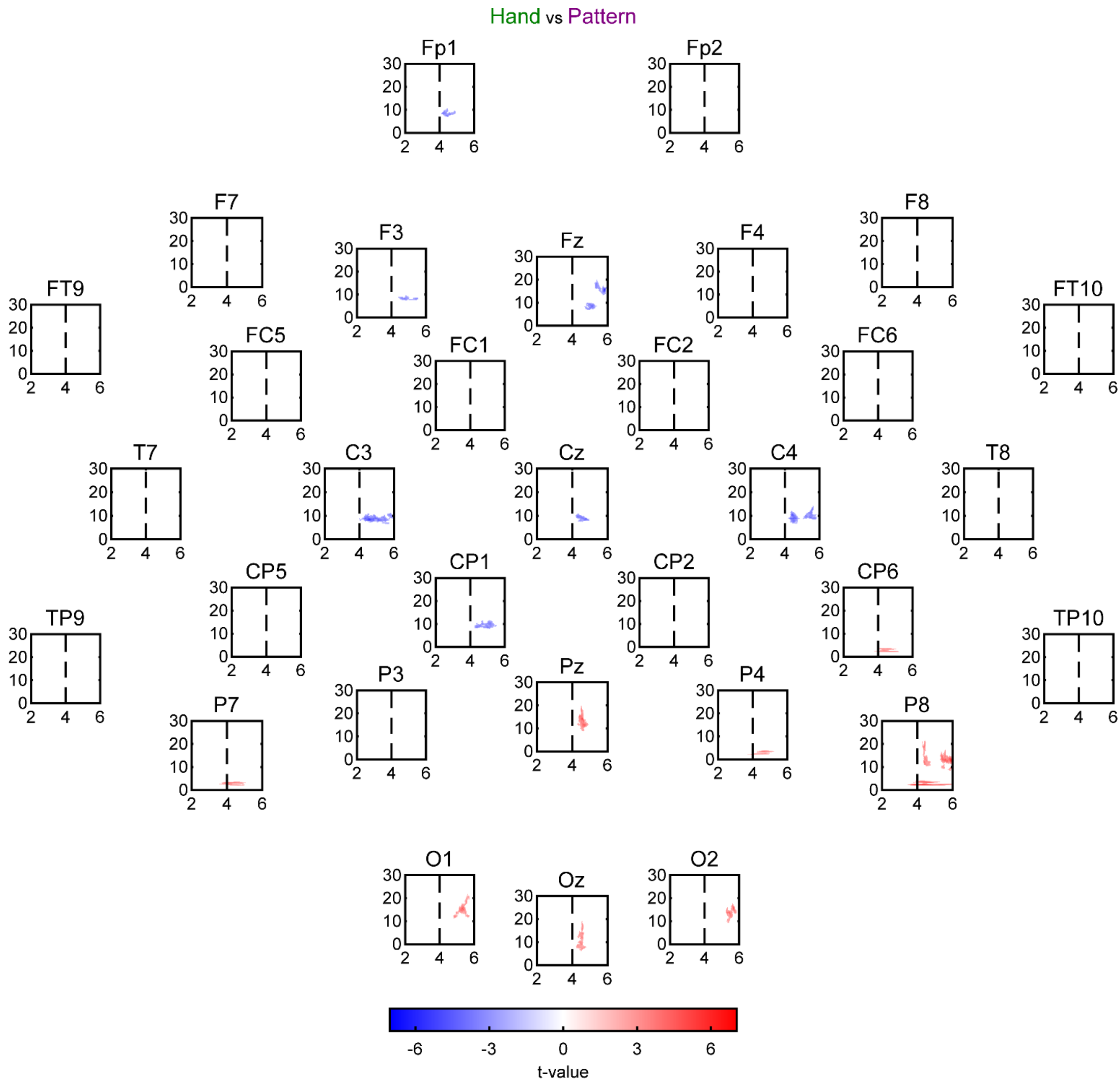


**Supplementary Figure 4. Time-frequency representations of Hand vs Pattern responses in 32 recording sites.** T-values of paired-samples t-tests comparing log power change values between Hand and Pattern videos across frequencies and time points in 32 recording sites. For all analyses, the sample size was 30 and the statistical significance was determined via cluster-correction. T-values of only significant clusters are shown. Colder colors indicate smaller log power change values for Hand, whereas warmer colors indicate smaller values for Pattern videos. Note that, during the initial 2 seconds, the video was static, and during the ladder 2 seconds, the video displayed motion.

### Supplementary Figure 5

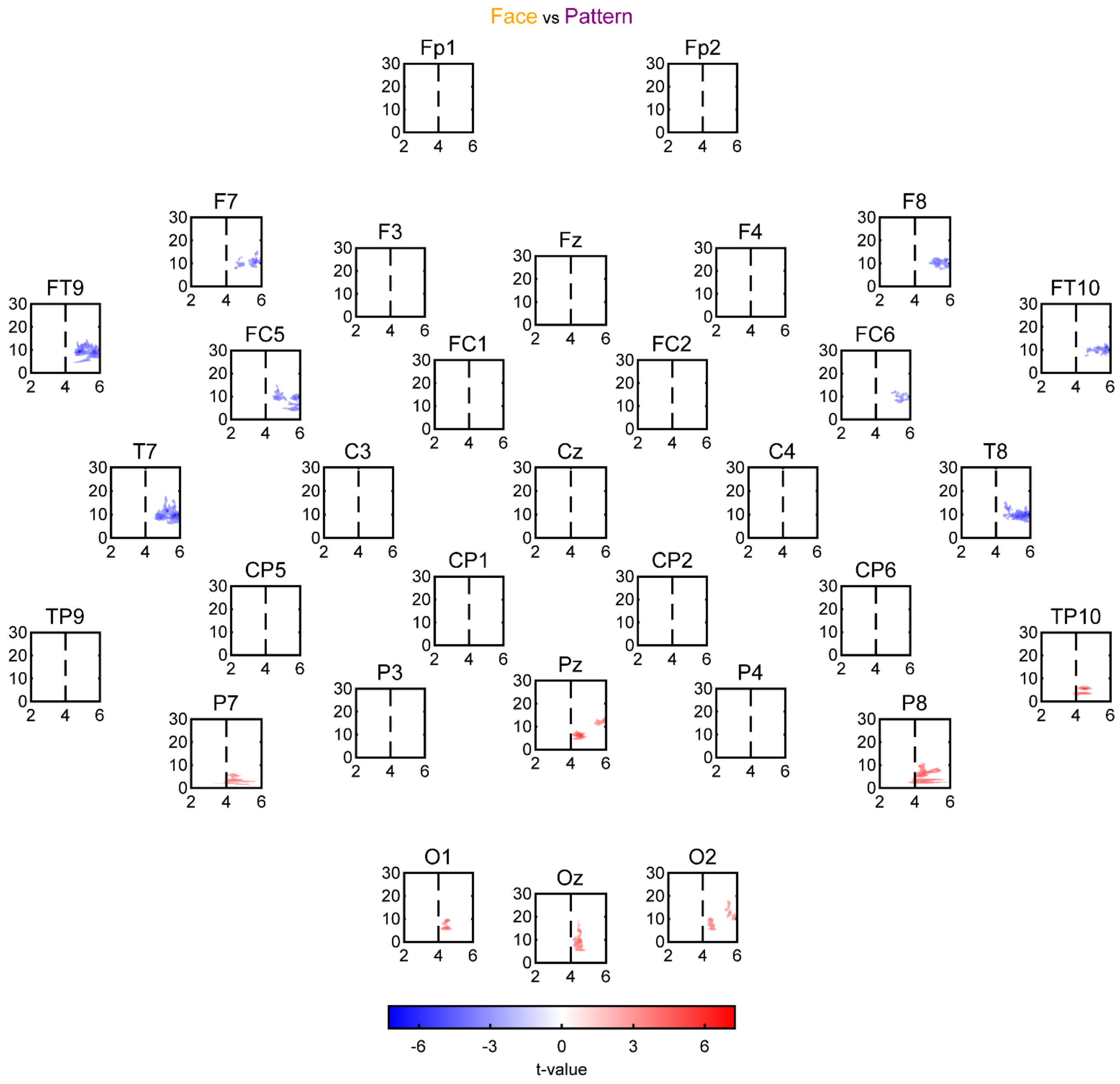


**Supplementary Figure 5. Time-frequency representations of Face vs Pattern responses in 32 recording sites.** T-values of paired-samples t-tests comparing log power change values between Face and Pattern videos across frequencies and time points in 32 recording sites. For all analyses, the sample size was 30 and the statistical significance was determined via cluster-correction. T-values of only significant clusters are shown. Colder colors indicate smaller log power change values for Face, whereas warmer colors indicate smaller values for Pattern videos. Note that, during the initial 2 seconds, the video was static, and during the ladder 2 seconds, the video displayed motion.

### Supplementary Figure 6

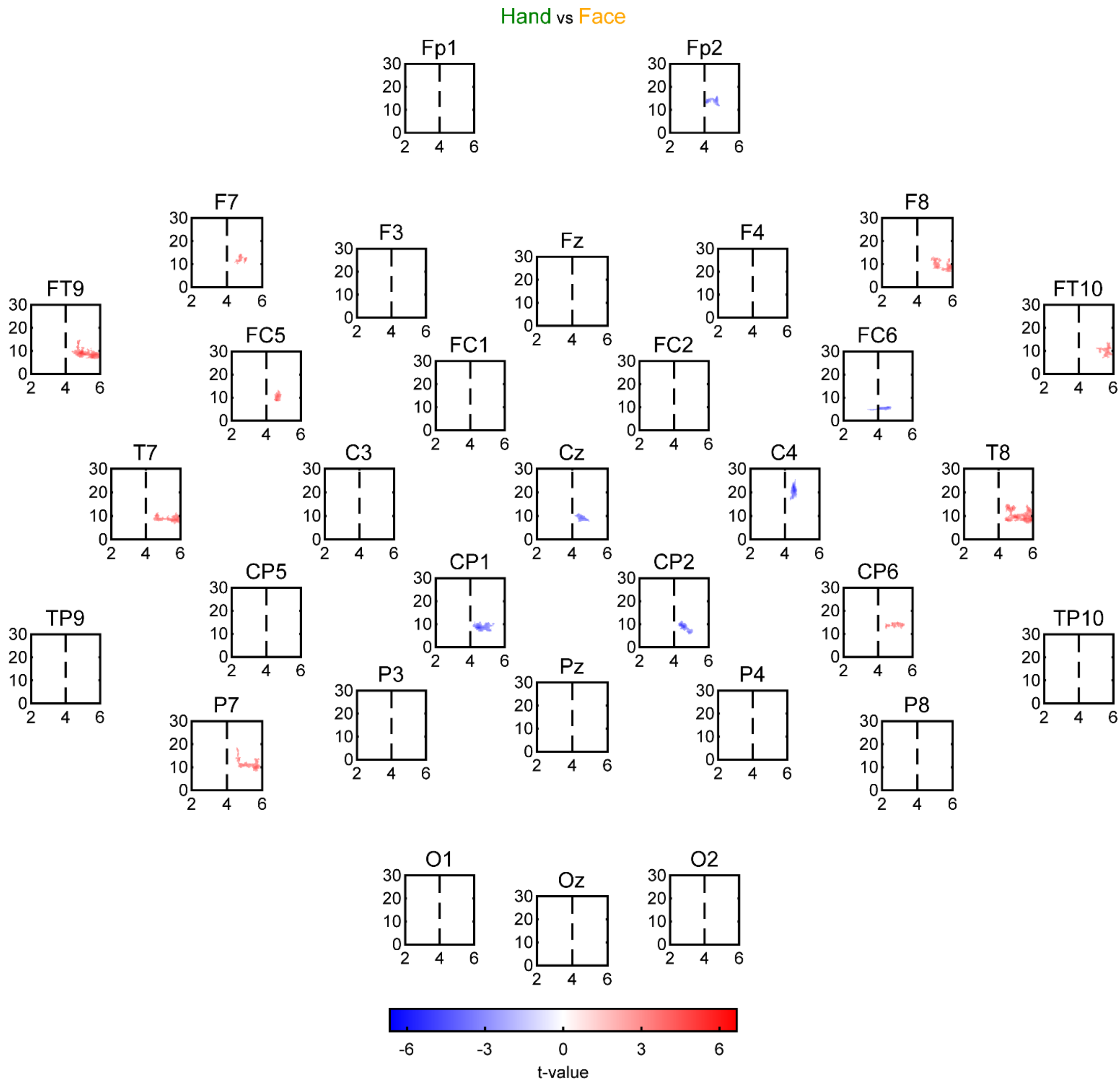


**Supplementary Figure 6. Time-frequency representations of Hand vs Face responses in 32 recording sites.** T-values of paired-samples t-tests comparing log power change values between Hand and Face videos across frequencies and time points in 32 recording sites. For all analyses, the sample size was 30 and the statistical significance was determined via cluster-correction. T-values of only significant clusters are shown. Colder colors indicate smaller log power change values for Hand, whereas warmer colors indicate smaller values for Face videos. Note that, during the initial 2 seconds, the video was static, and during the ladder 2 seconds, the video displayed motion.

### Supplementary Figure 7

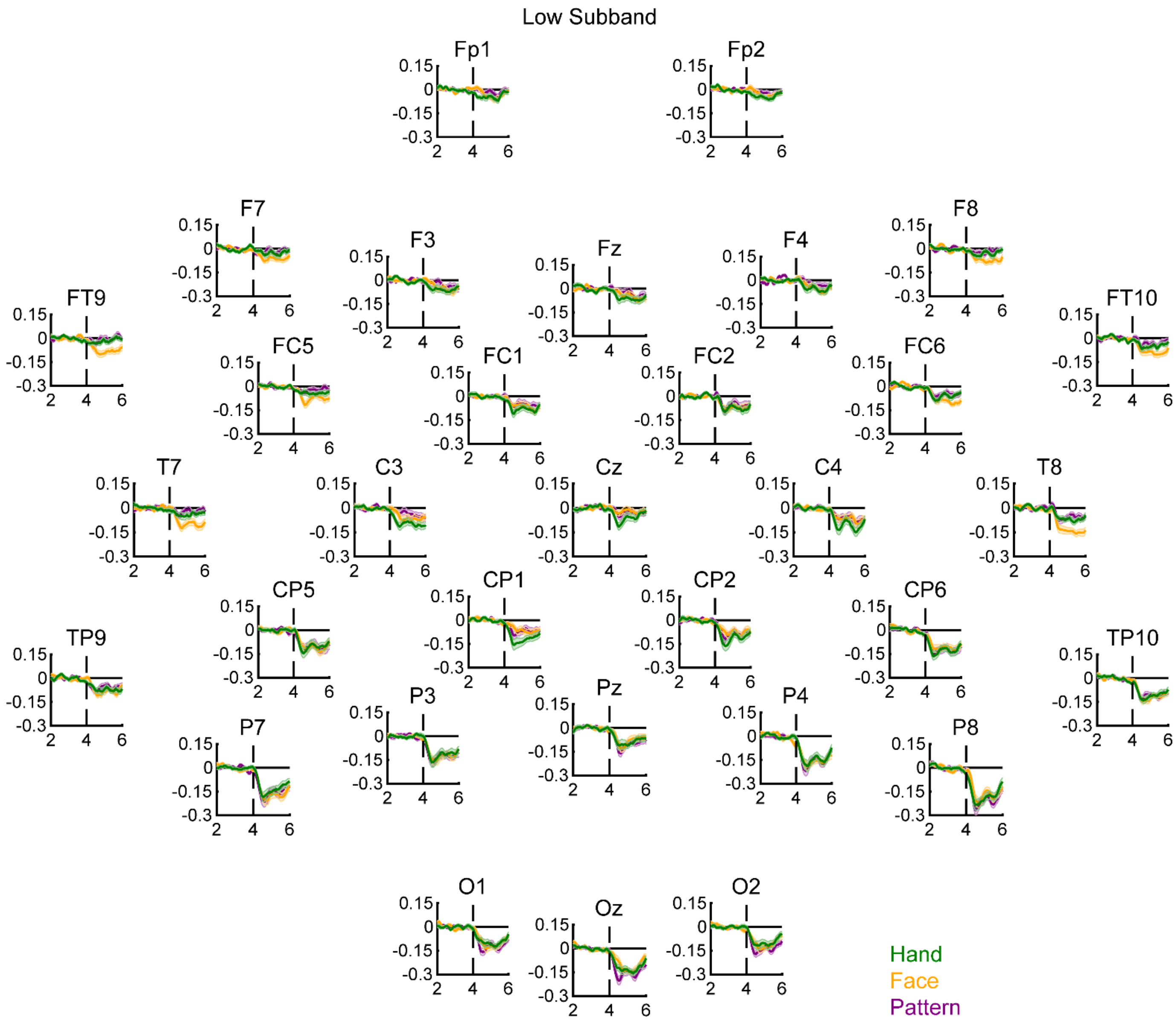


**Supplementary Figure 7. Temporal dynamics of Low subband responses in 32 recording sites.** Mean log power change values in the Low subband as a function of time for Hand, Face, and Pattern videos in 32 recording sites. Shadings depict SEMs. For all figures, the sample size was 30. Note that, during the initial 2 seconds, the video was static, and during the ladder 2 seconds, the video displayed motion.

### Supplementary Figure 8

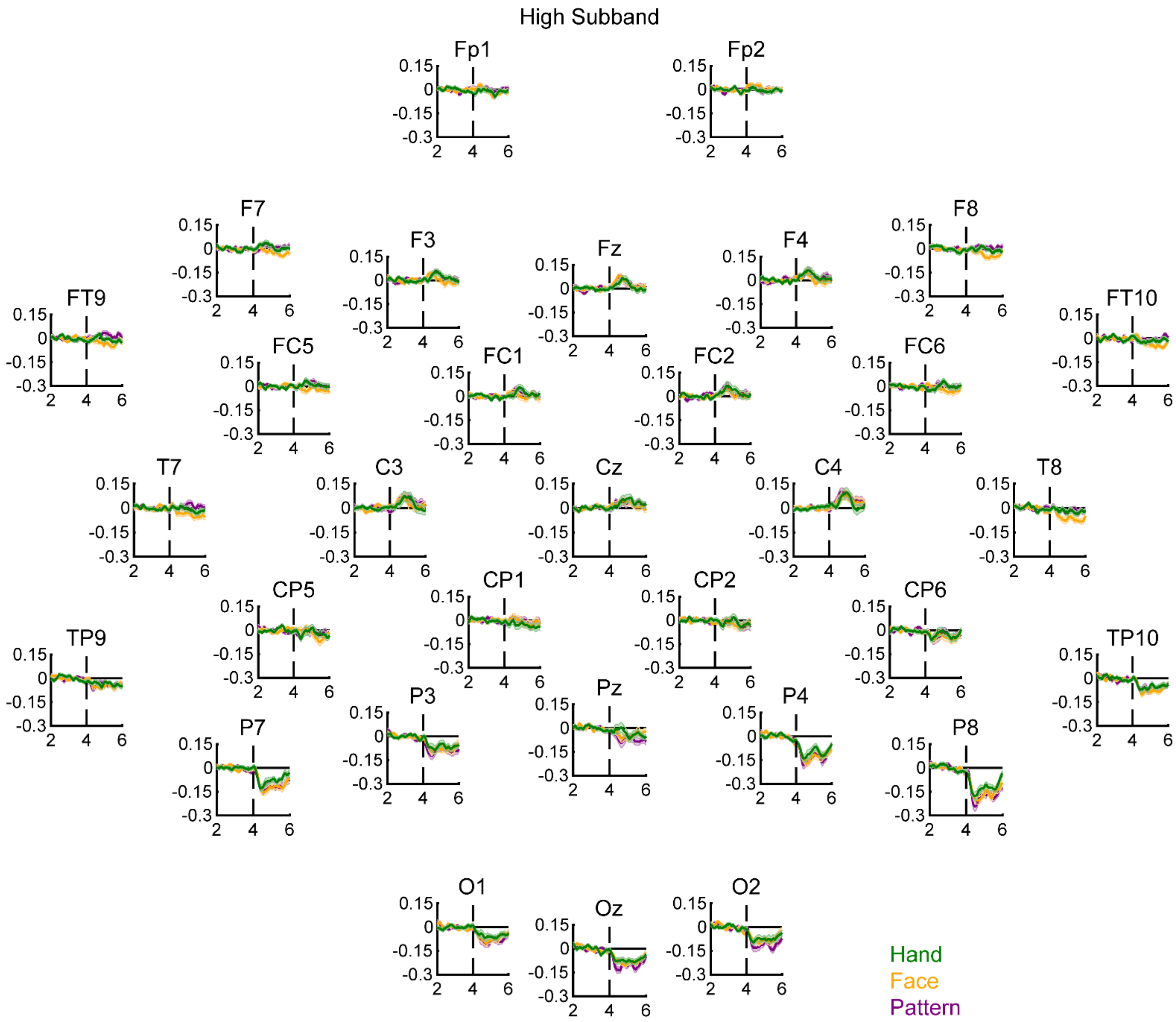


**Supplementary Figure 8. Temporal dynamics of High subband responses in 32 recording sites.** Mean log power change values in the High subband as a function of time for Hand, Face, and Pattern videos in 32 recording sites. Shadings depict SEMs. For all figures, the sample size was 30. Note that, during the initial 2 seconds, the video was static, and during the ladder 2 seconds, the video displayed motion.

### Supplementary Table 1

| Category | Action | Explanation |
|---|---|---|
| Hand | 1 | No movement |
| | 2 | Bending the middle finger from its base joint slightly towards the palm |
| | 3 | Bending the little finger from its base joint slightly towards the palm |
| | 4 | Moving the thumb laterally toward the index finger |
| | 5 | Moving the index and the middle fingers laterally toward each other |
| | 6 | Bending the index finger from its base joint strongly towards the palm |
| | 7 | Bending all the fingers from their middle joints slightly towards the palm |
| | 8 | Bending the middle and the ring fingers from their base joints strongly towards the palm |
| | 9 | Bending the index and the little fingers from their base joints strongly towards the palm |
| | 10 | Bending the middle, the ring, and the little fingers from their base joints strongly towards the palm |
| | 11 | Bending all the fingers from their middle joints strongly towards the palm |
| | 12 | Moving all fingers laterally toward each other |
| | 13 | Moving the thumb and the index finger so that their tips touch each other |
| Face | 1 | No movement |
| | 2 | Chewing while the mouth remains closed |
| | 3 | Blinking both eyes at the same time |
| | 4 | Lip puckering |
| | 5 | Protruding the lower lip while the mouth remains closed |
| | 6 | Raising both eyebrows at the same time |
| | 7 | Opening the mouth in the 'O' shape |
| | 8 | Pulling the nose up as in sniffing |
| | 9 | Opening the lips quickly to release air as in blowing |
| | 10 | Lifting the upper lip as in tooth-baring behavior while the teeth remain closed |
| | 11 | Puffing out cheeks |
| | 12 | Closing both eyes and pulling the mouth up at the same time while the mouth remains closed |
| | 13 | Raising both eyebrows and opening the mouth wide at the same time |

**Supplementary Table 1. Actions displayed in Hand and Face videos.** All actions were repeated twice at a once-per-second rate within the 2-second dynamic period of the videos. Hands and faces started all actions in their natural neutral state and returned to the same state after performing the actions. Actions are listed in ascending order according to their average motion energies with each stimulus category.

### Supplementary Table 2

| Subband | Period | Site | Stimulus | $t_{29}$ | $p$ | $BF_{10}$ |
|---|---|---|---|---|---|---|
| Low | Early | Central | Hand | **-4.581** | **$8 \times 10^{-5}$** | **313.724** |
| | | | Face | **-3.092** | **0.004** | **9.151** |
| | | | Pattern | -2.224 | 0.034 | 1.627 |
| | | Temporal | Hand | **-4.262** | **$2 \times 10^{-4}$** | **141.737** |
| | | | Face | **-5.139** | **$2 \times 10^{-5}$** | **1281.899** |
| | | | Pattern | -2.802 | 0.009 | 4.945 |
| | | Occipital | Hand | **-5.321** | **$10^{-5}$** | **2037.274** |
| | | | Face | **-6.000** | **$2 \times 10^{-6}$** | **$10^{4}$** |
| | | | Pattern | **-9.260** | **$4 \times 10^{-10}$** | **$3 \times 10^{7}$** |
| | Late | Central | Hand | **-5.765** | **$3 \times 10^{-6}$** | **6309.549** |
| | | | Face | **-3.608** | **0.001** | **29.356** |
| | | | Pattern | -2.834 | 0.008 | 5.278 |
| | | Temporal | Hand | **-4.389** | **$10^{-4}$** | **194.055** |
| | | | Face | **-7.019** | **$10^{-7}$** | **$2 \times 10^{5}$** |
| | | | Pattern | -2.619 | 0.014 | 3.419 |
| | | Occipital | Hand | **-6.012** | **$2 \times 10^{-6}$** | **$10^{4}$** |
| | | | Face | **-6.921** | **$10^{-7}$** | **$10^{5}$** |
| | | | Pattern | **-9.945** | **$8 \times 10^{-11}$** | **$10^{8}$** |
| High | Early | Central | Hand | 2.098 | 0.045 | 1.308 |
| | | | Face | 2.541 | 0.017 | 2.931 |
| | | | Pattern | **3.235** | **0.003** | **12.539** |
| | | Temporal | Hand | -0.625 | 0.537 | 0.233 |
| | | | Face | -2.719 | 0.011 | 4.174 |
| | | | Pattern | -0.389 | 0.700 | 0.209 |
| | | Occipital | Hand | **-4.338** | **$2 \times 10^{-4}$** | **171.274** |
| | | | Face | **-6.473** | **$4 \times 10^{-7}$** | **$4 \times 10^{4}$** |
| | | | Pattern | **-6.641** | **$3 \times 10^{-7}$** | **$6 \times 10^{4}$** |
| | Late | Central | Hand | 0.819 | 0.419 | 0.264 |
| | | | Face | 1.275 | 0.212 | 0.405 |
| | | | Pattern | 2.093 | 0.045 | 1.296 |
| | | Temporal | Hand | -2.150 | 0.040 | 1.429 |
| | | | Face | **-5.103** | **$2 \times 10^{-5}$** | **1170.235** |
| | | | Pattern | -0.112 | 0.912 | 0.196 |
| | | Occipital | Hand | **-4.443** | **$10^{-4}$** | **222.126** |
| | | | Face | **-4.811** | **$4 \times 10^{-5}$** | **559.429** |
| | | | Pattern | **-8.208** | **$5 \times 10^{-9}$** | **$3 \times 10^{6}$** |

**Supplementary Table 2. Statistics for one-sample t-tests comparing log power change values to zero.** Statistics were calculated for Low and High subbands in Early and Late periods for Hand, Face, and Pattern videos in Central, Temporal, and Occipital sites separately. For all analyses, the sample size was 30 and the significance level was Bonferroni-corrected at 0.05/9=0.006. Values significantly lower and higher than zero are highlighted in blue and red, respectively.

### Supplementary Table 3

| Subband | Period | Site | Stimulus Comparison | $t_{29}$ | $p$ | $BF_{10}$ |
|---|---|---|---|---|---|---|
| Low | Early | Central | Hand vs Pattern | **-5.367** | **$9\times10^{-6}$** | **2291.208** |
| | | | Face vs Pattern | -1.267 | 0.215 | 0.402 |
| | | | Hand vs Face | **-3.327** | **0.002** | **15.418** |
| | | Temporal | Hand vs Pattern | -1.062 | 0.297 | 0.325 |
| | | | Face vs Pattern | **-3.416** | **0.002** | **18.870** |
| | | | Hand vs Face | **3.212** | **0.003** | **11.899** |
| | | Occipital | Hand vs Pattern | 2.207 | 0.035 | 1.578 |
| | | | Face vs Pattern | 2.200 | 0.036 | 1.558 |
| | | | Hand vs Face | 0.329 | 0.745 | 0.204 |
| | Late | Central | Hand vs Pattern | **-4.022** | **$4\times10^{-4}$** | **78.894** |
| | | | Face vs Pattern | 0.899 | 0.376 | 0.282 |
| | | | Hand vs Face | **-3.149** | **0.004** | **10.360** |
| | | Temporal | Hand vs Pattern | -2.021 | 0.053 | 1.149 |
| | | | Face vs Pattern | **-6.848** | **$2\times10^{-7}$** | **$10^{5}$** |
| | | | Hand vs Face | **4.844** | **$4\times10^{-5}$** | **606.830** |
| | | Occipital | Hand vs Pattern | 1.886 | 0.069 | 0.925 |
| | | | Face vs Pattern | 1.291 | 0.207 | 0.412 |
| | | | Hand vs Face | 0.648 | 0.522 | 0.236 |
| High | Early | Central | Hand vs Pattern | -0.595 | 0.556 | 0.229 |
| | | | Face vs Pattern | -0.342 | 0.735 | 0.205 |
| | | | Hand vs Face | -0.304 | 0.764 | 0.203 |
| | | Temporal | Hand vs Pattern | -0.207 | 0.837 | 0.198 |
| | | | Face vs Pattern | -2.765 | 0.010 | 4.583 |
| | | | Hand vs Face | 2.358 | 0.025 | 2.074 |
| | | Occipital | Hand vs Pattern | 2.191 | 0.037 | 1.535 |
| | | | Face vs Pattern | 1.597 | 0.121 | 0.605 |
| | | | Hand vs Face | 1.301 | 0.203 | 0.418 |
| | Late | Central | Hand vs Pattern | -1.821 | 0.079 | 0.837 |
| | | | Face vs Pattern | -1.450 | 0.158 | 0.499 |
| | | | Hand vs Face | -0.498 | 0.622 | 0.218 |
| | | Temporal | Hand vs Pattern | -2.638 | 0.013 | 3.547 |
| | | | Face vs Pattern | **-7.420** | **$4\times10^{-8}$** | **$4\times10^{5}$** |
| | | | Hand vs Face | **3.397** | **0.002** | **18.042** |
| | | Occipital | Hand vs Pattern | 2.515 | 0.018 | 2.791 |
| | | | Face vs Pattern | 1.982 | 0.057 | 1.078 |
| | | | Hand vs Face | 0.485 | 0.631 | 0.217 |

**Supplementary Table 3. Statistics for paired-samples t-tests comparing log power change values between pairs of stimulus categories.** Statistics were calculated for Low and High subbands in Early and Late periods for Hand vs Pattern, Face vs Pattern, and Hand vs Face comparisons in Central, Temporal, and Occipital sites separately. For all analyses, the sample size was 30 and the significance level was Bonferroni-corrected at 0.05/9=0.006. Comparisons in which the first stimulus category in the stimulus comparison column is significantly lower or greater than the second stimulus category are highlighted with blue or red, respectively.

### Supplementary Table 4

| Subband | Period | Stimulus | Electrodes |
|---|---|---|---|
| Low | Early | Hand | Cz, C3, C4, CP1, CP2, CP5, CP6, Fz, F3, F8, FC1, FC2, FC5, FC6, Fp1, Fp2, FT10, Oz, O1, O2, Pz, P3, P4, P7, P8, T7, T8, TP9, TP10, F4, F7, FT9 |
| | | Face | C4, CP1, CP2, CP5, CP6, F7, F8, FC1, FC2, FC5, FC6, FT9, FT10, Oz, O1, O2, Pz, P3, P4, P7, P8, T7, T8, TP9, TP10, Cz, C3, Fz, F3, F4, Fp1, Fp2 |
| | | Pattern | CP1, CP2, CP5, CP6, FC6, Oz, O1, O2, Pz, P3, P4, P7, P8, T8, TP9, TP10, Cz, C3, C4, Fz, F3, F4, F7, F8, FC1, FC2, FC5, Fp1, Fp2, FT9, FT10, T7 |
| | Late | Hand | C3, C4, CP1, CP2, CP5, CP6, Fz, F3, F4, FC1, FC2, FC5, FC6, Fp1, Fp2, FT10, Oz, O1, O2, Pz, P3, P4, P7, P8, T8, TP9, TP10, Cz, F7, F8, FT9, T7 |
| | | Face | C3, C4, CP1, CP2, CP5, CP6, Fz, F3, F4, F7, F8, FC1, FC2, FC5, FC6, Fp1, Fp2, FT9, FT10, Oz, O1, O2, Pz, P3, P4, P7, P8, T7, T8, TP9, TP10, Cz |
| | | Pattern | C4, CP1, CP2, CP5, CP6, FC1, FC6, Oz, O1, O2, Pz, P3, P4, P7, P8, T8, TP9, TP10, Cz, C3, Fz, F3, F4, F7, F8, FC2, FC5, Fp1, Fp2, FT9, FT10, T7 |
| High | Early | Hand | Oz, O2, P4, P7, P8, TP9, TP10, Cz, C3, C4, CP1, CP2, CP5, CP6, Fz, F3, F4, F7, F8, FC1, FC2, FC5, FC6, Fp1, Fp2, FT9, FT10, O1, Pz, P3, T7, T8 |
| | | Face | Oz, O1, O2, P3, P4, P7, P8, TP9, TP10, Fz, Cz, C3, C4, CP1, CP2, CP5, CP6, F3, F4, F7, F8, FC1, FC2, FC5, FC6, Fp1, Fp2, FT9, FT10, Pz, T7, T8 |
| | | Pattern | Oz, O1, O2, Pz, P3, P4, P7, P8, TP9, TP10, C4, Fz, F3, Cz, C3, C4, CP1, CP2, CP5, CP6, F4, F7, F8, FC1, FC2, FC5, FC6, Fp1, Fp2, FT9, FT10, T7, T8 |
| | Late | Hand | Oz, O1, O2, P3, P4, P7, P8, TP9, TP10, Cz, C3, C4, CP1, CP2, CP5, CP6, Fz, F3, F4, F7, F8, FC1, FC2, FC5, FC6, Fp1, Fp2, FT9, FT10, Pz, T7, T8 |
| | | Face | F7, F8, FT9, FT10, Fp1, Oz, O1, O2, P3, P4, P7, P8, T7, T8, TP9, TP10, Fz, Cz, C3, C4, CP1, CP2, CP5, CP6, F3, F4, FC1, FC2, FC5, FC6, Fp2, Pz |
| | | Pattern | Oz, O1, O2, Pz, P3, P4, P7, P8, TP9, TP10, C4, Fz, F3, Cz, C3, C4, CP1, CP2, CP5, CP6, F4, F7, F8, FC1, FC2, FC5, FC6, Fp1, Fp2, FT9, FT10, T7, T8 |

**Supplementary Table 4. Statistical significance of one-sample t-tests comparing log power change values to zero in 32 recording sites.** Statistics were calculated for Low and High subbands in Early and Late periods for Hand, Face, and Pattern videos separately in 32 recording sites. For all analyses, the sample size was 30 and the significance level was corrected for false discovery rate at a q level of 0.05. Sites with values significantly lower and higher than zero are highlighted in blue and red, respectively. Not highlighted sites did not reveal statistical significance.

### Supplementary Table 5

| Subband | Period | Stimulus Comparison | Electrodes |
|---|---|---|---|
| Low | Early | Hand vs Pattern | **Cz, C3, C4, Fp1,** CP1, CP2, CP5, CP6, Fz, F3, F4, F7, F8, FC1, FC2, FC5, FC6, Fp2, FT9, FT10, Oz, O1, O2, Pz, P3, P4, P7, P8, T7, T8, TP9, TP10 |
| | | Face vs Pattern | Cz, C3, C4, CP1, CP2, CP5, CP6, Fz, F3, F4, F7, F8, FC1, FC2, FC5, FC6, Fp1, Fp2, FT9, FT10, Oz, O1, O2, Pz, P3, P4, P7, P8, T7, T8, TP9, TP10 |
| | | Hand vs Face | Cz, C3, C4, CP1, CP2, CP5, CP6, Fz, F3, F4, F7, F8, FC1, FC2, FC5, FC6, Fp1, Fp2, FT9, FT10, Oz, O1, O2, Pz, P3, P4, P7, P8, T7, T8, TP9, TP10 |
| | Late | Hand vs Pattern | Cz, C3, C4, CP1, CP2, CP5, CP6, Fz, F3, F4, F7, F8, FC1, FC2, FC5, FC6, Fp1, Fp2, FT9, FT10, Oz, O1, O2, Pz, P3, P4, P7, P8, T7, T8, TP9, TP10 |
| | | Face vs Pattern | **F7, F8, FC5, FC6, FT9, FT10, T7, T8,** Cz, C3, C4, CP1, CP2, CP5, CP6, Fz, F3, F4, FC1, FC2, Fp1, Fp2, Oz, O1, O2, Pz, P3, P4, P7, P8, TP9, TP10 |
| | | Hand vs Face | **F8, FT9, T7, T8,** Cz, C3, C4, CP1, CP2, CP5, CP6, Fz, F3, F4, F7, FC1, FC2, FC5, FC6, Fp1, Fp2, FT10, Oz, O1, O2, Pz, P3, P4, P7, P8, TP9, TP10 |
| High | Early | Hand vs Pattern | Cz, C3, C4, CP1, CP2, CP5, CP6, Fz, F3, F4, F7, F8, FC1, FC2, FC5, FC6, Fp1, Fp2, FT9, FT10, Oz, O1, O2, Pz, P3, P4, P7, P8, T7, T8, TP9, TP10 |
| | | Face vs Pattern | Cz, C3, C4, CP1, CP2, CP5, CP6, Fz, F3, F4, F7, F8, FC1, FC2, FC5, FC6, Fp1, Fp2, FT9, FT10, Oz, O1, O2, Pz, P3, P4, P7, P8, T7, T8, TP9, TP10 |
| | | Hand vs Face | Cz, C3, C4, CP1, CP2, CP5, CP6, Fz, F3, F4, F7, F8, FC1, FC2, FC5, FC6, Fp1, Fp2, FT9, FT10, Oz, O1, O2, Pz, P3, P4, P7, P8, T7, T8, TP9, TP10 |
| | Late | Hand vs Pattern | Cz, C3, C4, CP1, CP2, CP5, CP6, Fz, F3, F4, F7, F8, FC1, FC2, FC5, FC6, Fp1, Fp2, FT9, FT10, Oz, O1, O2, Pz, P3, P4, P7, P8, T7, T8, TP9, TP10 |
| | | Face vs Pattern | **F7, F8, FT9, FT10, T7, T8,** Cz, C3, C4, CP1, CP2, CP5, CP6, Fz, F3, F4, FC1, FC2, FC5, FC6, Fp1, Fp2, Oz, O1, O2, Pz, P3, P4, P7, P8, TP9, TP10 |
| | | Hand vs Face | Cz, C3, C4, CP1, CP2, CP5, CP6, Fz, F3, F4, F7, F8, FC1, FC2, FC5, FC6, Fp1, Fp2, FT9, FT10, Oz, O1, O2, Pz, P3, P4, P7, P8, T7, T8, TP9, TP10 |

**Supplementary Table 5. Statistical significance of paired-sample t-tests comparing log power change values between pairs of stimulus categories in 32 recording sites.** Statistics were calculated for Low and High subbands in Early and Late periods for Hand vs Pattern, Face vs Pattern, and Hand vs Face videos separately in 32 recording sites. For all analyses, the sample size was 30 and the significance level was corrected for false discovery rate at a q level of 0.05. For each comparison, sites in which the first stimulus category in the stimulus comparison column is significantly lower or greater than the second stimulus category are highlighted with blue or red, respectively. Not highlighted sites did not reveal statistical significance.

### Supplementary Table 6

| Subband | Period | Site | Stimulus | $t_{60}$ | $p$ | $BF_{10}$ |
|---|---|---|---|---|---|---|
| Low | Early | Central | Hand-No-Object | **-4.072** | **$10^{-4}$** | **160.623** |
| | | | Hand-Object | **-2.943** | **0.005** | **6.829** |
| | | | Kaleidoscope | -2.551 | 0.013 | 2.722 |
| | | Occipital | Hand-No-Object | **-4.917** | **$7x10^{-6}$** | **2485.716** |
| | | | Hand-Object | **-6.349** | **$3x10^{-8}$** | **$4x10^{5}$** |
| | | | Kaleidoscope | **-8.114** | **$3x10^{-11}$** | **$3x10^{8}$** |
| | Late | Central | Hand-No-Object | **-3.001** | **0.004** | **7.891** |
| | | | Hand-Object | **-2.854** | **0.006** | **5.499** |
| | | | Kaleidoscope | **-3.846** | **$3x10^{-4}$** | **80.907** |
| | | Occipital | Hand-No-Object | **-5.600** | **$6x10^{-7}$** | **$3x10^{4}$** |
| | | | Hand-Object | **-5.902** | **$2x10^{-7}$** | **$8x10^{4}$** |
| | | | Kaleidoscope | **-8.525** | **$6x10^{-12}$** | **$10^{9}$** |
| High | Early | Central | Hand-No-Object | 1.224 | 0.226 | 0.285 |
| | | | Hand-Object | 1.932 | 0.058 | 0.796 |
| | | | Kaleidoscope | **3.025** | **0.004** | **8.393** |
| | | Occipital | Hand-No-Object | **-5.397** | **$10^{-6}$** | **$10^{4}$** |
| | | | Hand-Object | **-5.572** | **$6x10^{-7}$** | **$2x10^{4}$** |
| | | | Kaleidoscope | **-7.800** | **$10^{-10}$** | **$9x10^{7}$** |
| | Late | Central | Hand-No-Object | 2.579 | 0.012 | 2.898 |
| | | | Hand-Object | 2.452 | 0.017 | 2.196 |
| | | | Kaleidoscope | 2.435 | 0.018 | 2.114 |
| | | Occipital | Hand-No-Object | **-4.781** | **$10^{-5}$** | **1574.756** |
| | | | Hand-Object | **-5.704** | **$4x10^{-7}$** | **$4x10^{4}$** |
| | | | Kaleidoscope | **-7.862** | **$8x10^{-11}$** | **$10^{8}$** |

**Supplementary Table 6. Statistics for one-sample t-tests comparing log power change values to zero in the Hobson and Bishop (2016) dataset.** Statistics were calculated for Low and High subbands in the Early and Late periods for Hand-No-Object, Hand-Object, and Kaleidoscope videos in Central and Occipital sites separately. For all analyses, the sample size was 61 and the significance level was Bonferroni-corrected at 0.05/6=0.008. Values significantly lower and higher than zero are highlighted in blue and red, respectively.

### Supplementary Table 7

| Subband | Period | Site | Stimulus Comparison | $t_{60}$ | $p$ | $BF_{10}$ |
|---|---|---|---|---|---|---|
| Low | Early | Central | HNO vs Kaleidoscope | -2.072 | 0.043 | 1.026 |
| | | | HO vs Kaleidoscope | -0.963 | 0.339 | 0.218 |
| | | | HNO vs HO | -1.031 | 0.307 | 0.232 |
| | | Occipital | HNO vs Kaleidoscope | **4.529** | **$3x10^{-5}$** | **684.703** |
| | | | HO vs Kaleidoscope | **3.495** | **$9x10^{-4}$** | **29.362** |
| | | | HNO vs HO | 1.324 | 0.190 | 0.321 |
| | Late | Central | HNO vs Kaleidoscope | 0.342 | 0.734 | 0.148 |
| | | | HO vs Kaleidoscope | 0.918 | 0.362 | 0.209 |
| | | | HNO vs HO | -0.593 | 0.556 | 0.166 |
| | | Occipital | HNO vs Kaleidoscope | **5.001** | **$5x10^{-6}$** | **3312.472** |
| | | | HO vs Kaleidoscope | **4.774** | **$10^{-5}$** | **1535.484** |
| | | | HNO vs HO | -0.187 | 0.852 | 0.143 |
| High | Early | Central | HNO vs Kaleidoscope | -1.442 | 0.155 | 0.374 |
| | | | HO vs Kaleidoscope | -0.820 | 0.415 | 0.193 |
| | | | HNO vs HO | -0.670 | 0.505 | 0.174 |
| | | Occipital | HNO vs Kaleidoscope | **3.223** | **0.002** | **14.000** |
| | | | HO vs Kaleidoscope | **2.904** | **$2x10^{-4}$** | **96.158** |
| | | | HNO vs HO | -0.973 | 0.335 | 0.220 |
| | Late | Central | HNO vs Kaleidoscope | 0.434 | 0.666 | 0.153 |
| | | | HO vs Kaleidoscope | -0.107 | 0.915 | 0.141 |
| | | | HNO vs HO | 0.682 | 0.498 | 0.175 |
| | | Occipital | HNO vs Kaleidoscope | **4.819** | **$10^{-5}$** | **1785.282** |
| | | | HO vs Kaleidoscope | **4.616** | **$2x10^{-5}$** | **909.676** |
| | | | HNO vs HO | -0.427 | 0.671 | 0.153 |

**Supplementary Table 7. Statistics for paired-samples t-tests comparing log power change values between pairs of stimulus categories in the Hobson and Bishop (2016) dataset.** Statistics were calculated for Low and High subbands in Early and Late periods for Hand-No-Object (HNO) vs Kaleidoscope, Hand-Object (HO) vs Pattern, and HNO vs HO comparisons in Central and Occipital sites separately. For all analyses, the sample size was 30 and the significance level was Bonferroni-corrected at 0.05/6=0.008. Comparisons in which the first stimulus category in the stimulus comparison column is significantly greater than the second stimulus category are highlighted in red.

### Supplementary Table 8

| Subband | Period | Site | Stimulus | $t_{90}$ | $p$ | $BF_{10}$ |
|---|---|---|---|---|---|---|
| Low | Early | Central | Hand & HNO | **-5.959** | **$5 \times 10^{-8}$** | **$3 \times 10^{5}$** |
| | | | Pattern & Kaleidoscope | **-3.387** | **0.001** | **22.044** |
| | | Occipital | Hand & HNO | **-6.983** | **$5 \times 10^{-10}$** | **$2 \times 10^{7}$** |
| | | | Pattern & Kaleidoscope | **-11.326** | **$5 \times 10^{-19}$** | **$10^{16}$** |
| | Late | Central | Hand & HNO | **-5.234** | **$10^{-6}$** | **$10^{4}$** |
| | | | Pattern & Kaleidoscope | **-4.800** | **$6 \times 10^{-6}$** | **2514.646** |
| | | Occipital | Hand & HNO | **-7.834** | **$9 \times 10^{-12}$** | **$10^{9}$** |
| | | | Pattern & Kaleidoscope | **-11.579** | **$2 \times 10^{-19}$** | **$3 \times 10^{16}$** |
| High | Early | Central | Hand & HNO | 2.226 | 0.029 | 1.205 |
| | | | Pattern & Kaleidoscope | **4.323** | **$4 \times 10^{-5}$** | **448.340** |
| | | Occipital | Hand & HNO | **-6.922** | **$6 \times 10^{-10}$** | **$2 \times 10^{7}$** |
| | | | Pattern & Kaleidoscope | **-10.171** | **$10^{-16}$** | **$5 \times 10^{13}$** |
| | Late | Central | Hand & HNO | **2.616** | **0.010** | **2.853** |
| | | | Pattern & Kaleidoscope | **3.189** | **0.002** | **12.516** |
| | | Occipital | Hand & HNO | **-6.357** | **$8 \times 10^{-9}$** | **$10^{6}$** |
| | | | Pattern & Kaleidoscope | **-10.233** | **$9 \times 10^{-17}$** | **$7 \times 10^{13}$** |

**Supplementary Table 8. Statistics for one-sample t-tests comparing log power change values to zero in the combined dataset of the present study and the Hobson and Bishop (2016) study.** Statistics were calculated for Low and High subbands in the Early and Late periods by combining Hand and Hand-No-Object (HNO) together and Pattern and Kaleidoscope videos together separately in Central and Occipital sites. For all analyses, the combined sample size was 91 and the significance level was Bonferroni-corrected at 0.05/4=0.013. Values significantly lower and higher than zero are highlighted in blue and red, respectively.

### Supplementary Table 9

| Subband | Period | Site | Stimulus Comparison | $t_{90}$ | $p$ | $BF_{10}$ |
|---|---|---|---|---|---|---|
| Low | Early | Central | Hand & HNO vs Pattern & Kaleidoscope | **-4.062** | **$10^{-4}$** | **183.908** |
| | | Occipital | Hand & HNO vs Pattern & Kaleidoscope | **4.994** | **$3x10^{-6}$** | **5236.281** |
| | Late | Central | Hand & HNO vs Pattern & Kaleidoscope | -1.490 | 0.140 | 0.336 |
| | | Occipital | Hand & HNO vs Pattern & Kaleidoscope | **5.192** | **$10^{-6}$** | **$10^{4}$** |
| High | Early | Central | Hand & HNO vs Pattern & Kaleidoscope | -1.560 | 0.122 | 0.372 |
| | | Occipital | Hand & HNO vs Pattern & Kaleidoscope | **3.910** | **$2x10^{-4}$** | **111.343** |
| | Late | Central | Hand & HNO vs Pattern & Kaleidoscope | -0.422 | 0.674 | 0.126 |
| | | Occipital | Hand & HNO vs Pattern & Kaleidoscope | **5.408** | **$5x10^{-7}$** | **$3x10^{4}$** |

**Supplementary Table 9. Statistics for paired-samples t-tests comparing log power change values between pairs of stimulus categories in the combined dataset of the present study and the Hobson and Bishop (2016) study.** Statistics were calculated for Low and High subbands in the Early and Late periods by combining Hand and Hand-No-Object (HNO) together and Pattern and Kaleidoscope videos together separately in Central and Occipital sites. For all analyses, the combined sample size was 91 and the significance level was Bonferroni-corrected at 0.05/2=0.025. Comparisons in which the first stimulus category in the stimulus comparison column is lower or greater than the second stimulus category are highlighted with blue or red, respectively.

### Supplementary Table 10

| Subband | Period | Site | Stimulus | $t_{29}$ | $p$ | $BF_{10}$ |
|---|---|---|---|---|---|---|
| Low | Early | Central | Hand | -2.590 | 0.015 | 3.228 |
| | | | Face | 0.691 | 0.495 | 0.242 |
| | | | Pattern | 1.518 | 0.140 | 0.545 |
| | | Temporal | Hand | -0.785 | 0.439 | 0.258 |
| | | | Face | -1.301 | 0.203 | 0.417 |
| | | | Pattern | -0.866 | 0.393 | 0.274 |
| | | Occipital | Hand | **-5.747** | **$3x10^{-6}$** | **6026.249** |
| | | | Face | -1.129 | 0.268 | 0.347 |
| | | | Pattern | 0.768 | 0.449 | 0.255 |
| | Late | Central | Hand | -1.915 | 0.065 | 0.968 |
| | | | Face | -0.960 | 0.345 | 0.296 |
| | | | Pattern | 0.812 | 0.423 | 0.263 |
| | | Temporal | Hand | -1.602 | 0.120 | 0.610 |
| | | | Face | -0.964 | 0.343 | 0.297 |
| | | | Pattern | -0.957 | 0.346 | 0.296 |
| | | Occipital | Hand | **-3.336** | **0.002** | **15.705** |
| | | | Face | -0.057 | 0.955 | 0.195 |
| | | | Pattern | 0.013 | 0.990 | 0.194 |
| High | Early | Central | Hand | -1.876 | 0.071 | 0.911 |
| | | | Face | 0.102 | 0.919 | 0.195 |
| | | | Pattern | 0.933 | 0.359 | 0.289 |
| | | Temporal | Hand | 1.265 | 0.216 | 0.401 |
| | | | Face | -0.547 | 0.589 | 0.223 |
| | | | Pattern | -0.850 | 0.402 | 0.271 |
| | | Occipital | Hand | -2.790 | 0.009 | 4.823 |
| | | | Face | -2.617 | 0.014 | 3.401 |
| | | | Pattern | 1.273 | 0.213 | 0.405 |
| | Late | Central | Hand | 1.635 | 0.113 | 0.638 |
| | | | Face | -1.342 | 0.190 | 0.438 |
| | | | Pattern | 0.035 | 0.972 | 0.195 |
| | | Temporal | Hand | -0.633 | 0.532 | 0.234 |
| | | | Face | -2.825 | 0.008 | 5.184 |
| | | | Pattern | 0.040 | 0.968 | 0.195 |
| | | Occipital | Hand | -1.850 | 0.074 | 0.875 |
| | | | Face | -1.717 | 0.097 | 0.717 |
| | | | Pattern | 1.127 | 0.269 | 0.346 |

**Supplementary Table 10. Statistics for one-sample t-tests comparing correlation coefficients between perceived motion ratings in the validation study and log power change values to zero.** Statistics were calculated for Low and High subbands in Early and Late periods for Hand, Face, and Pattern videos in Central, Temporal, and Occipital sites separately. Correlations coefficients were calculated with the average motion ratings of videos in the validation study. For all analyses, the sample size was 30 and the significance level was Bonferroni-corrected at 0.05/9=0.006. Values significantly lower than zero are highlighted in blue.

### Supplementary Table 11

| Subband | Period | Site | Stimulus | $t_{29}$ | $p$ | $BF_{10}$ |
|---|---|---|---|---|---|---|
| Low | Early | Central | Hand | -2.798 | 0.009 | 4.902 |
| | | | Face | 0.760 | 0.453 | 0.254 |
| | | | Pattern | 1.639 | 0.114 | 0.633 |
| | | Temporal | Hand | -0.655 | 0.518 | 0.237 |
| | | | Face | -0.634 | 0.531 | 0.234 |
| | | | Pattern | -0.894 | 0.379 | 0.280 |
| | | Occipital | Hand | **-4.720** | **$6x10^{-5}$** | **444.709** |
| | | | Face | -2.340 | 0.026 | 2.007 |
| | | | Pattern | 0.516 | 0.610 | 0.220 |
| | Late | Central | Hand | -2.241 | 0.033 | 1.676 |
| | | | Face | -1.271 | 0.214 | 0.403 |
| | | | Pattern | 1.062 | 0.297 | 0.325 |
| | | Temporal | Hand | -2.129 | 0.042 | 1.378 |
| | | | Face | -0.905 | 0.373 | 0.283 |
| | | | Pattern | -0.807 | 0.426 | 0.262 |
| | | Occipital | Hand | **-3.402** | **0.002** | **18.263** |
| | | | Face | -0.908 | 0.371 | 0.284 |
| | | | Pattern | -0.131 | 0.896 | 0.196 |
| High | Early | Central | Hand | **-3.665** | **$10^{-5}$** | **33.570** |
| | | | Face | -0.003 | 0.998 | 0.194 |
| | | | Pattern | 0.751 | 0.459 | 0.252 |
| | | Temporal | Hand | 1.310 | 0.201 | 0.422 |
| | | | Face | 0.891 | 0.381 | 0.280 |
| | | | Pattern | -0.745 | 0.462 | 0.251 |
| | | Occipital | Hand | -1.530 | 0.137 | 0.554 |
| | | | Face | -2.724 | 0.011 | 4.219 |
| | | | Pattern | 1.165 | 0.253 | 0.360 |
| | Late | Central | Hand | 1.542 | 0.134 | 0.562 |
| | | | Face | -1.155 | 0.257 | 0.356 |
| | | | Pattern | 0.269 | 0.790 | 0.201 |
| | | Temporal | Hand | 0.156 | 0.877 | 0.197 |
| | | | Face | -2.277 | 0.030 | 1.789 |
| | | | Pattern | 0.172 | 0.865 | 0.197 |
| | | Occipital | Hand | -1.564 | 0.129 | 0.579 |
| | | | Face | -1.582 | 0.125 | 0.593 |
| | | | Pattern | 1.048 | 0.303 | 0.321 |

**Supplementary Table 11. Statistics for one-sample t-tests comparing correlation coefficients between motion energy estimations and log power change values to zero.** Statistics were calculated for Low and High subbands in Early and Late periods for Hand, Face, and Pattern videos in Central, Temporal, and Occipital sites separately. Correlations coefficients were calculated with the time-averaged motion energy estimations of videos. For all analyses, the sample size was 30 and the significance level was Bonferroni-corrected at 0.05/9=0.006. Values significantly lower than zero are highlighted in blue.

### Supplementary Table 12

| Subband | Period | Site | Stimulus | $t_{29}$ | $p$ | $BF_{10}$ |
|---|---|---|---|---|---|---|
| Low | Early | Central | Hand | -2.011 | 0.054 | 1.131 |
| | | | Face | -0.499 | 0.622 | 0.218 |
| | | | Pattern | 1.684 | 0.103 | 0.683 |
| | | Temporal | Hand | -0.245 | 0.808 | 0.200 |
| | | | Face | -2.087 | 0.046 | 1.283 |
| | | | Pattern | 0.147 | 0.884 | 0.196 |
| | | Occipital | Hand | **-4.080** | **$3x10^{-4}$** | **90.699** |
| | | | Face | -1.726 | 0.095 | 0.726 |
| | | | Pattern | 1.694 | 0.101 | 0.694 |
| | Late | Central | Hand | -2.341 | 0.026 | 2.010 |
| | | | Face | -1.300 | 0.204 | 0.417 |
| | | | Pattern | 0.030 | 0.976 | 0.194 |
| | | Temporal | Hand | -1.541 | 0.134 | 0.562 |
| | | | Face | -2.507 | 0.018 | 2.744 |
| | | | Pattern | 0.952 | 0.349 | 0.294 |
| | | Occipital | Hand | -2.192 | 0.037 | 1.537 |
| | | | Face | -0.708 | 0.485 | 0.245 |
| | | | Pattern | 1.257 | 0.219 | 0.397 |
| High | Early | Central | Hand | -1.402 | 0.171 | 0.470 |
| | | | Face | 0.006 | 0.995 | 0.194 |
| | | | Pattern | 1.525 | 0.138 | 0.550 |
| | | Temporal | Hand | 1.323 | 0.196 | 0.428 |
| | | | Face | -0.660 | 0.514 | 0.238 |
| | | | Pattern | 1.395 | 0.174 | 0.467 |
| | | Occipital | Hand | -2.955 | 0.006 | 6.806 |
| | | | Face | **-3.134** | **0.004** | **10.032** |
| | | | Pattern | 2.037 | 0.051 | 1.180 |
| | Late | Central | Hand | 0.304 | 0.763 | 0.203 |
| | | | Face | -0.433 | 0.668 | 0.212 |
| | | | Pattern | 0.440 | 0.663 | 0.213 |
| | | Temporal | Hand | -0.641 | 0.527 | 0.235 |
| | | | Face | -2.910 | 0.007 | 6.198 |
| | | | Pattern | 2.538 | 0.017 | 2.916 |
| | | Occipital | Hand | -1.493 | 0.146 | 0.527 |
| | | | Face | -2.117 | 0.043 | 1.352 |
| | | | Pattern | 2.392 | 0.023 | 2.206 |

**Supplementary Table 12. Statistics for one-sample t-tests comparing correlation coefficients between participants' own motion ratings and log power change values to zero.** Statistics were calculated for Low and High subbands in Early and Late periods for Hand, Face, and Pattern videos in Central, Temporal, and Occipital sites separately. For each participant, correlation coefficients were calculated with the motion ratings of that participant. Since only a quarter of all trials were rated by each participant, the specific rating given to each unique video was also used in the other non-rated trials with different versions (original, horizontally flipped) and other repetitions of the same video during the calculation of these correlation coefficients. For all analyses, the sample size was 30 and the significance level was Bonferroni-corrected at 0.05/9=0.006. Values significantly lower than zero are highlighted in blue.

### Supplementary Table 13

| Subband | Period | Site | Stimulus Pair | $r_{28}$ | $p$ | $BF_{10}$ |
|---|---|---|---|---|---|---|
| Low | Early | Central | Hand - Pattern | **0.870** | **$4x10^{-10}$** | **$3x10^{7}$** |
| | | | Face - Pattern | **0.838** | **$8x10^{-9}$** | **$2x10^{6}$** |
| | | | Hand - Face | **0.794** | **$2x10^{-7}$** | **$10^{5}$** |
| | | Temporal | Hand - Pattern | **0.654** | **$9x10^{-5}$** | **344.468** |
| | | | Face - Pattern | 0.488 | 0.006 | 8.050 |
| | | | Hand - Face | **0.632** | **$2x10^{-4}$** | **182.828** |
| | | Occipital | Hand - Pattern | **0.629** | **$2x10^{-4}$** | **154.127** |
| | | | Face - Pattern | **0.709** | **$10^{-5}$** | **2137.678** |
| | | | Hand - Face | **0.609** | **$4x10^{-4}$** | **98.847** |
| | Late | Central | Hand - Pattern | **0.812** | **$5x10^{-8}$** | **$3x10^{5}$** |
| | | | Face - Pattern | **0.812** | **$5x10^{-8}$** | **$3x10^{5}$** |
| | | | Hand - Face | **0.798** | **$10^{-7}$** | **$10^{5}$** |
| | | Temporal | Hand - Pattern | **0.716** | **$9x10^{-6}$** | **2790.238** |
| | | | Face - Pattern | **0.676** | **$4x10^{-5}$** | **673.267** |
| | | | Hand - Face | **0.583** | **$7x10^{-4}$** | **53.298** |
| | | Occipital | Hand - Pattern | **0.524** | **0.003** | **15.261** |
| | | | Face - Pattern | 0.401 | 0.028 | 2.274 |
| | | | Hand - Face | **0.738** | **$3x10^{-6}$** | **6749.070** |
| High | Early | Central | Hand - Pattern | **0.789** | **$2x10^{-7}$** | **$8x10^{4}$** |
| | | | Face - Pattern | **0.847** | **$4x10^{-9}$** | **$4x10^{6}$** |
| | | | Hand - Face | **0.760** | **$10^{-6}$** | **$2x10^{4}$** |
| | | Temporal | Hand - Pattern | **0.548** | **0.002** | **27.798** |
| | | | Face - Pattern | **0.521** | **0.003** | **14.469** |
| | | | Hand - Face | 0.381 | 0.038 | 1.772 |
| | | Occipital | Hand - Pattern | 0.475 | 0.008 | 6.495 |
| | | | Face - Pattern | **0.633** | **$2x10^{-4}$** | **185.676** |
| | | | Hand - Face | **0.680** | **$4x10^{-5}$** | **779.146** |
| | Late | Central | Hand - Pattern | **0.816** | **$4x10^{-8}$** | **$4x10^{5}$** |
| | | | Face - Pattern | **0.847** | **$4x10^{-9}$** | **$3x10^{6}$** |
| | | | Hand - Face | **0.752** | **$2x10^{-6}$** | **$10^{4}$** |
| | | Temporal | Hand - Pattern | **0.671** | **$5x10^{-5}$** | **571.590** |
| | | | Face - Pattern | **0.738** | **$3x10^{-6}$** | **6725.085** |
| | | | Hand - Face | 0.476 | 0.008 | 6.625 |
| | | Occipital | Hand - Pattern | **0.512** | **0.004** | **12.206** |
| | | | Face - Pattern | 0.381 | 0.038 | 1.788 |
| | | | Hand - Face | **0.818** | **$3x10^{-8}$** | **$5x10^{5}$** |

**Supplementary Table 13. Statistics for correlations between the log power change values of pairs of stimulus categories.** Statistics were calculated for Low and High subbands in Early and Late periods for Hand and Pattern, Face and Pattern, and Hand and Face stimulus pairs in Central, Temporal, and Occipital sites separately. For all analyses, the sample size was 30 and the significance level was Bonferroni-corrected at 0.05/9=0.006. Significantly positive correlations are highlighted in red.

### Supplementary Table 14

| Subband | Period | Site | Stimulus | $r_{28}$ | *p* | $BF_{10}$ |
|---|---|---|---|---|---|---|
| Low | Early | Central | Hand | 0.233 | 0.216 | 0.472 |
| | | | Face | 0.319 | 0.085 | 0.932 |
| | | | Pattern | 0.199 | 0.291 | 0.387 |
| | | Temporal | Hand | 0.058 | 0.760 | 0.237 |
| | | | Face | 0.032 | 0.868 | 0.230 |
| | | | Pattern | 0.114 | 0.547 | 0.270 |
| | | Occipital | Hand | 0.294 | 0.114 | 0.746 |
| | | | Face | 0.291 | 0.246 | 0.432 |
| | | | Pattern | -0.125 | 0.510 | 0.279 |
| | Late | Central | Hand | 0.107 | 0.574 | 0.264 |
| | | | Face | 0.197 | 0.296 | 0.382 |
| | | | Pattern | 0.077 | 0.686 | 0.245 |
| | | Temporal | Hand | 0.039 | 0.838 | 0.231 |
| | | | Face | -0.035 | 0.854 | 0.231 |
| | | | Pattern | 0.263 | 0.160 | 0.582 |
| | | Occipital | Hand | 0.149 | 0.434 | 0.304 |
| | | | Face | 0.151 | 0.425 | 0.308 |
| | | | Pattern | -0.135 | 0.478 | 0.289 |
| High | Early | Central | Hand | -0.056 | 0.767 | 0.237 |
| | | | Face | 0.022 | 0.907 | 0.228 |
| | | | Pattern | 0.097 | 0.611 | 0.257 |
| | | Temporal | Hand | -0.072 | 0.705 | 0.243 |
| | | | Face | 0.024 | 0.902 | 0.229 |
| | | | Pattern | 0.150 | 0.430 | 0.306 |
| | | Occipital | Hand | 0.124 | 0.514 | 0.278 |
| | | | Face | -0.171 | 0.366 | 0.335 |
| | | | Pattern | -0.158 | 0.403 | 0.317 |
| | Late | Central | Hand | -0.055 | 0.775 | 0.236 |
| | | | Face | 0.101 | 0.595 | 0.260 |
| | | | Pattern | 0.027 | 0.889 | 0.229 |
| | | Temporal | Hand | 0.058 | 0.759 | 0.237 |
| | | | Face | -0.103 | 0.587 | 0.261 |
| | | | Pattern | 0.141 | 0.458 | 0.295 |
| | | Occipital | Hand | 0.124 | 0.515 | 0.278 |
| | | | Face | 0.121 | 0.524 | 0.276 |
| | | | Pattern | -0.318 | 0.087 | 0.918 |

**Supplementary Table 14. Statistics for correlations between the log power change values and Interpersonal Reactivity Index scores.** Statistics were calculated for Low and High subbands in the Early and Late periods for Hand, Face, and Pattern, videos in Central, Temporal, and Occipital sites separately. For all analyses, the sample size was 30 and the significance level was Bonferroni-corrected at 0.05/9=0.006.

### Supplementary Table 15

| | Correlation Coefficient | | | | Number of Trials | | |
|---|---|---|---|---|---|---|---|
| **Participant** | **Hand** | **Face** | **Pattern** | **All** | **Hand** | **Face** | **Pattern** |
| 1 | 0.479 | 0.400 | 0.106 | 0.227 | 46 | 68 | 44 |
| 2 | 0.204 | 0.555 | 0.222 | 0.340 | 32 | 65 | 37 |
| 3 | 0.880 | 0.703 | 0.758 | 0.753 | 67 | 90 | 80 |
| 4 | 0.843 | 0.592 | 0.517 | 0.672 | 72 | 78 | 76 |
| 8 | 0.365 | 0.376 | 0.352 | 0.390 | 82 | 89 | 87 |
| 10 | 0.216 | 0.357 | 0.291 | 0.240 | 67 | 86 | 71 |
| 11 | 0.602 | 0.585 | 0.528 | 0.491 | 75 | 82 | 83 |
| 12 | 0.335 | 0.241 | 0.515 | 0.402 | 53 | 56 | 50 |
| 13 | 0.404 | 0.319 | 0.497 | 0.414 | 43 | 36 | 51 |
| 14 | 0.385 | 0.232 | 0.293 | 0.332 | 79 | 91 | 82 |
| 15 | 0.357 | 0.349 | 0.463 | 0.348 | 49 | 56 | 61 |
| 16 | 0.377 | 0.408 | 0.390 | 0.426 | 75 | 82 | 74 |
| 17 | 0.373 | 0.445 | 0.404 | 0.430 | 72 | 70 | 68 |
| 18 | 0.676 | 0.630 | 0.501 | 0.643 | 73 | 89 | 66 |
| 19 | 0.700 | 0.747 | 0.706 | 0.712 | 102 | 103 | 101 |
| 21 | 0.871 | 0.741 | 0.647 | 0.711 | 100 | 99 | 97 |
| 23 | 0.137 | 0.313 | 0.312 | 0.276 | 45 | 44 | 35 |
| 24 | 0.525 | 0.446 | 0.668 | 0.437 | 76 | 82 | 79 |
| 25 | 0.646 | 0.736 | 0.418 | 0.594 | 58 | 65 | 45 |
| 26 | 0.608 | 0.813 | 0.623 | 0.646 | 91 | 95 | 96 |
| 27 | 0.351 | 0.319 | 0.250 | 0.288 | 30 | 33 | 35 |
| 28 | 0.694 | 0.741 | 0.724 | 0.670 | 89 | 97 | 87 |
| 29 | 0.411 | 0.532 | 0.171 | 0.330 | 59 | 69 | 52 |
| 31 | 0.624 | 0.406 | 0.415 | 0.474 | 25 | 39 | 29 |
| 32 | 0.722 | 0.750 | 0.513 | 0.680 | 93 | 90 | 96 |
| 34 | 0.158 | 0.300 | 0.228 | 0.287 | 30 | 46 | 27 |
| 36 | 0.406 | 0.186 | 0.354 | 0.327 | 84 | 90 | 92 |
| 37 | 0.786 | 0.776 | 0.758 | 0.792 | 101 | 96 | 99 |
| 39 | 0.816 | 0.718 | 0.315 | 0.541 | 52 | 66 | 52 |
| 41 | 0.672 | 0.702 | 0.634 | 0.688 | 96 | 96 | 91 |
| **Mean (SD)** | **0.521 (0.215)** | **0.514 (0.192)** | **0.453 (0.179)** | **0.485 (0.171)** | **67 (23)** | **75 (20)** | **68 (23)** |

**Supplementary Table 15. Motion rating correlations and numbers of included trials in EEG analyses.** For each participant, correlation coefficients were calculated between the average motion ratings in the validation study and that participant's own motion ratings. Since only a quarter of all trials were rated, correlation coefficients in the 'All' column, which combined all videos, were estimated at 312/4=78 trials, whereas the stimulus-specific correlation coefficients were each estimated at 78/3=26 trials. Thus, the critical r values in two-tailed tests with alpha levels of 0.05 were 0.223 and 0.361 for the combined and each stimulus-specific correlation coefficient, respectively. The number of trials included in EEG analyses is 312/3=104 for each stimulus category.
